# AARS2 ameliorates myocardial ischemia *via* fine-tuning PKM2-mediated metabolism

**DOI:** 10.1101/2024.06.04.597368

**Authors:** Zongwang Zhang, Lixia Zheng, Yang Chen, Yuanyuan Chen, Junjie Hou, Chenglu Xiao, Xiaojun Zhu, Shi-Min Zhao, Jing-Wei Xiong

## Abstract

AARS2, an alanyl-tRNA synthase, is essential for protein translation, but its function in mouse hearts is not fully addressed. Here, we found that cardiomyocyte-specific deletion of mouse AARS2 exhibited evident cardiomyopathy with impaired cardiac function, notable cardiac fibrosis and cardiomyocyte apoptosis. Cardiomyocyte-specific AARS2 overexpression in mice improved cardiac function and reduced cardiac fibrosis after myocardial infarction (MI), without affecting cardiomyocyte proliferation and coronary angiogenesis. Mechanistically, AARS2 overexpression suppressed cardiomyocyte apoptosis and mitochondrial reactive oxide species production, and changed cellular metabolism from oxidative phosphorylation toward glycolysis in cardiomyocytes, thus leading to cardiomyocyte survival from ischemia and hypoxia stress. Ribo-Seq revealed that AARS2 overexpression increased pyruvate kinase M2 (PKM2) protein translation and the ratio of PKM2 dimers to tetramers that promote glycolysis. Additionally, PKM2 activator TEPP-46 reversed cardiomyocyte apoptosis and cardiac fibrosis caused by AARS2 deficiency. Thus, this study demonstrates that AARS2 plays an essential role in protecting cardiomyocytes from ischemic pressure *via* fine-tuning PKM2-mediated energy metabolism, and presents a novel cardiac protective AARS2-PKM2 signaling during the pathogenesis of MI.

## Introduction

Cardiovascular disease (CVD) remains the leading cause of death worldwide (Mendis, Davis, & Norrving, 2015). Myocardial infarction (MI), a major type of CVDs, is considered an epidemic, affecting ∼1 to 2 percent of adults and posing a serious threat to human health and life (Thygesen et al., 2018). MI is due to coronary atherosclerosis and coronary heart disease, and MI patients can develop arrhythmia, shock, heart failure, and even death in a short time in severe cases (Frangogiannis, 2015; Thygesen et al., 2018). MI results in massive loss of cardiomyocytes and abnormal cardiac remodeling with severe inflammation and fibrosis (Q. Zhang et al., 2022). However, it is incompletely understood how to prevent massive cardiomyocyte death and inhibit cardiac fibrosis in time to achieve cardiac repair. Therefore, the main goals of cardiovascular research are to improve the adverse consequences of MI by improving immune inflammation and cardiac remodeling, cardiomyocyte protection, cardiomyocyte regeneration, and avoiding the expansion of fibrosis and scars.

As the heart ages, the reduced integrity and increased risk of heart diseases such as myocardial ischemia frequently take place, ultimately leading to heart failure (Strait & Lakatta, 2012). Due to limited regenerative potential of mammalian cardiomyocytes, massive death of cardiomyocytes leads to a decrease in their numbers, causing myocardial remodeling, hypertrophy, and impaired cardiac function after MI (Whelan, Kaplinskiy, & Kitsis, 2010). Over the years, the most effective approach to reducing acute myocardial ischemic injury is to salvage surviving cardiomyocytes. During ischemia, oxygen supply is reduced, leading to a decrease in ATP generation (Kalogeris, Baines, Krenz, & Korthuis, 2012). The significant reduction in ATP levels during ischemia is associated with the development of irreversible changes in cardiomyocytes because the cells deplete their energy reserves and cannot maintain internal homeostasis (Saraste, 1999). To compensate for the reduced oxidative phosphorylation (OXPHOS) during ischemia, cardiomyocytes increase glycolysis to maintain homeostasis. Under extreme stress conditions, such as MI, cardiomyocytes experience toxic levels of Ca^2+^ and reactive oxygen species (ROS). Mitochondrial electron transport chain (ETC) uncoupling leads to excessive ROS production, resulting in lipid and protein oxidation, widespread cell damage, and initiation of cardiomyocyte death (Penna, Mancardi, Rastaldo, & Pagliaro, 2009). The detrimental effects of ROS include DNA and RNA damage, lipid peroxidation, protein oxidation, and functional impairments (Schieber & Chandel, 2014). Mitochondria play a vital role in various metabolic pathways (Sissler, González-Serrano, & Westhof, 2017). Mitochondria protein mutations, which are involved in mitochondrial metabolism and directly affect the central functions of mitochondria, have been extensively studied for a long time (Gorman et al., 2016). On the other hand, mitochondrial dysfunction and mutations in different components of the mitochondrial translation machinery may indirectly impact ATP production (Pearce, Nezich, & Spinazzola, 2013). These includes mitochondrial aminoacyl-tRNA synthetases (mt-AARSs).

Mt-AARSs are a group of nuclei-coding enzymes that ensure the accurate translation of the genetic code by binding 20 amino acids to their homologous tRNA molecules (Meyer-Schuman & Antonellis, 2017; Ognjenović & Simonović, 2018; Sissler et al., 2017). Cytoplasmic AARS provides amino acid-tRNA binding for protein translation, and the corresponding mt-AARS are introduced into the mitochondrial matrix, fulfilling their typical role of transporting amino acids for tRNA molecules encoded by their mitochondrial genomes (mt-tRNA). The accuracy of mitochondrial protein synthesis depends on the coordination of nuclear encoded mt-AARS and mitochondrial DNA-encoded tRNA (Ognjenović & Simonović, 2018). Mitochondrial dysfunction is known to preferentially affect tissues with high energy requirements, especially the brain, muscles, and heart. It is widely recognized that nearly all nuclear genes associated with mt-AARS are considered pathogenic genes responsible for mitochondrial diseases (Euro et al., 2015). Typically, mutations in the mitochondrial alanyl-tRNA synthetase (AARS2) gene cause infantile-onset cardiomyopathy and white matter brain disease from childhood to adulthood (Nielsen et al., 2020). AARS2 is the gene encoding mitochondrial alanyl-tRNA synthetase, which is responsible for connecting specific tRNA with its cognate amino acid, alanine (X. Zhang et al., 2022). Recent studies have shown that AARS2 contains proline-hydroxylated conserved amino acid sequences, and its stability is enhanced under hypoxic conditions (Mao et al., 2024). Mutations in AARS2 can cause mitochondrial dysfunction (Dallabona et al., 2014; Vasilescu et al., 2018), leading to a reduction in cellular energy production (Fine, Nemeth, Kaufman, & Fatemi, 2019). However, the precise mechanisms by which AARS2 deficiency contributes to the development of cardiomyopathy in patients are not fully understood, and the potential of AARS2 as a therapeutic target for heart-related diseases remains to be explored.

One of the nodal points of energy metabolism is pyruvate kinase that catalyzes the conversion of phosphoenolpyruvate (PEP) to pyruvate, the last step of glycolysis (Méndez-Lucas et al., 2017). Mammalian pyruvate kinases (PKs) are encoded by two genes (PKLR and PKM), and each can generate two isoforms, respectively (PKL and PKR; PKM1 and PKM2) (Wong, Ojo, Yan, & Tang, 2015). Pyruvate kinase muscle isozyme 1 (PKM1) and pyruvate kinase muscle isozyme 2 (PKM2) are alternative splicing products of the PKM gene and express in various tissues (Israelsen & Vander Heiden, 2015). PKM1 is found in some differentiated adult tissues, such as heart, muscle, and brain. PKM2 is widely expressed and a predominant isoform in many adult cell types, including kidney tubular cells, intestinal epithelial cells, and lung epithelial cells (Dayton, Gocheva, et al., 2016). PKM1 has constitutively high catalytic activity, whereas PKM2 enzyme activity is subject to complex allosteric regulation (Ikeda, Tanaka, & Noguchi, 1997), which allows cells to switch between glycolysis and biosynthesis (Tang et al., 2023). PKM2 was recently found to protect cardiomyocytes from ischemia and promote cardiomyocyte proliferation (Magadum et al., 2020; Ni et al., 2022; Wu et al., 2021). However, it remains to be addressed how PKM2 protein and mRNA are regulated during pathogenesis of MI.

We have previously reported that AARS2 binds lactate to modulate metabolic proteins in ischemia hearts and skeletal muscles (Du et al., 2022; Mao et al., 2024). However, we have very limited knowledge on AARS2 function in the heart. And our previous experiments demonstrated that AARS2 overexpression had no significant effect on pyruvate dehydrogenase PDHA1 lactylation in cardiomyocytes. Here, we investigated AARS2 function under ischemic and hypoxic stress by employing both loss-of-function and gain-of-function of AARS2 in mice and neonatal rat cardiomyocytes (NRCMs). And we deciphered the mechanistic insights of AARS2 into energy metabolism by using variety of cell biology, biochemistry, mass spectrometry, and transcriptomics technologies. This work not only reveals a novel cardiac protection signaling AARS2-PKM2, but also identifies therapeutic targets and small molecules for cardiomyopathy and MI.

## Results

### Deleting AARS2 in adult cardiomyocytes causes heart failure

2020; Vasilescu et al., 2018), but the underlying mechanisms remain unclear. By comparing expression pattern of AARS2 before and after mouse MI, we found that both AARS2 proteins and mRNA decreased in the hearts after MI (Figure 1A–B), suggesting that AARS2 might be involved in the pathological progression of MI. To elucidate AARS2 function in adult cardiomyocytes, we crossed AARS2^floxed/floxed^ (AARS2^fl/fl^) mice with α-MHC-MerCreMer mice to achieve cardiomyocyte-specific knockout of AARS2 after tamoxifen treatment (Figure 1C). We used AARS2^fl/fl^ mice as control group, and α-MHC-MerCreMer; AARS2^fl/fl^ mice as cKO group (Figure 1–figure supplement 1A). As expected, AARS2 conditioned knockout (AARS2^cKO^) hearts had lower levels of AARS2 than that of AARS2^fl/fl^ hearts (Figure 1D), while the levels of AARS2 were comparable in the liver, lung and skeletal muscle between AARS2^fl/fl^ and cKO groups (Figure 1–figure supplement 1B). The results suggest that we succeed in achieving myocardial-specific deletion of AARS2 in adult cardiomyocytes.

**Figure 1.**
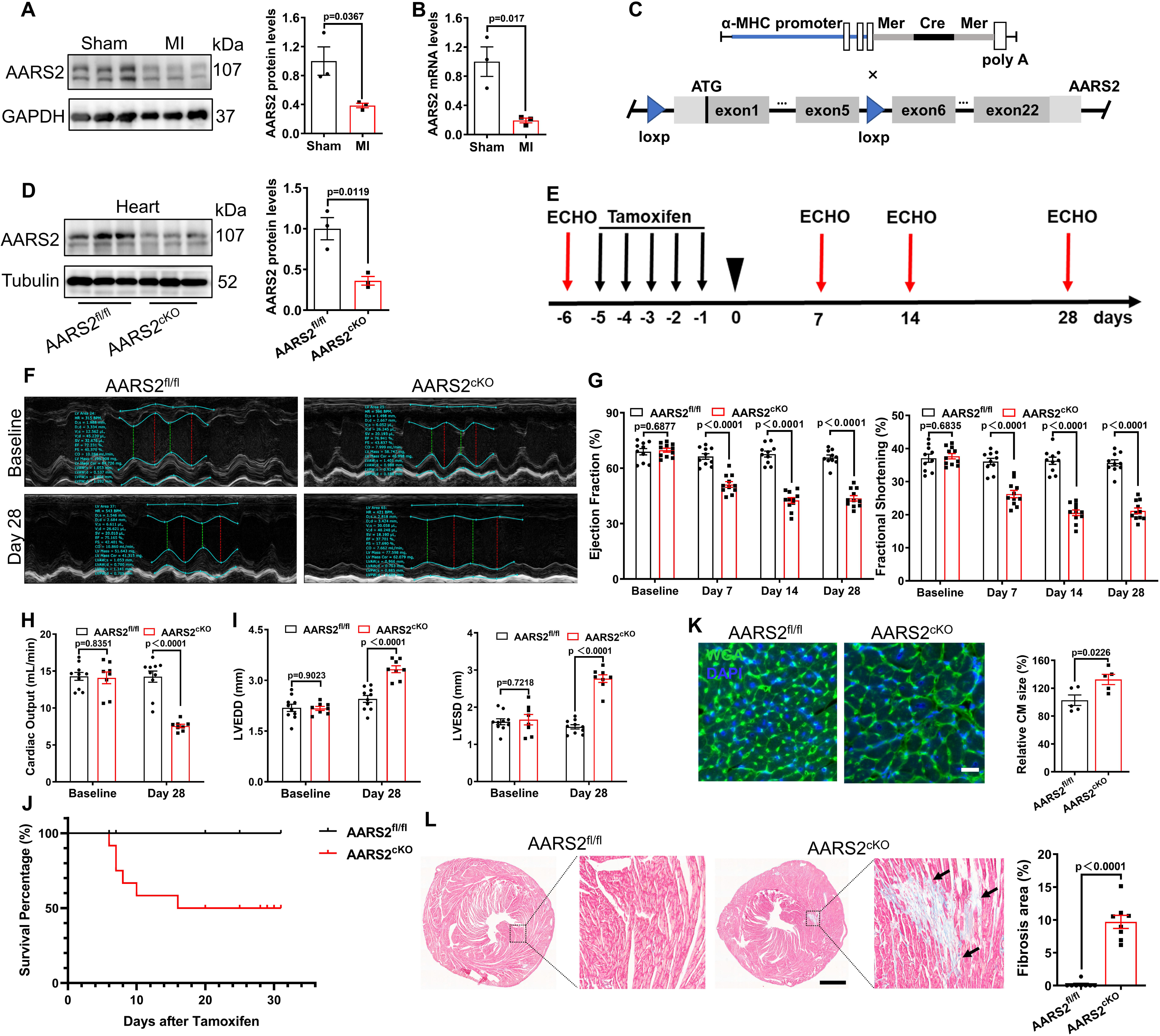
Cardiomyocyte-specific knockout of AARS2 leads to cardiac dysfunction and fibrosis in mice. (A–B) Western blot and qRT-PCR analysis showing reduced expression of AARS2 proteins (A) or mRNA (B) of 3-day MI hearts compared with sham hearts (n = 3). (C) Construction diagram of α-MHC-MerCreMer (upper) and AARS2^fl/fl^ mice (lower). (D) Western blots showing reduced AARS2 proteins in AARS2^cKO^ hearts compared with AARS2^fl/fl^ hearts (n = 3). (E) Schematic timelines of tamoxifen treatment and echocardiography (ECHO). (F) Representative M-mode tracings of ECHO in control and cKO hearts before and after tamoxifen treatment. (G) Ejection fraction (EF) and fractional shortening (FS) of AARS2^fl/fl^ and AARS2^cKO^ hearts at different time points after tamoxifen induction (n = 10–11). (H) Cardiac output of AARS2^fl/fl^ and AARS2^cKO^ mice at 28 days after tamoxifen induction (n = 8–10). (I) Left ventricular end-diastolic diameter (LVEDD) and left ventricular end-systolic diameter (LVESD) in AARS2^fl/fl^ and AARS2^cKO^ hearts at 28 days after tamoxifen induction (n = 8–10). (J) Survival percentage of AARS2^fl/fl^ and AARS2^cKO^ mice at 28 days after tamoxifen treatment (n = 8–10). (K) WGA immunofluorescence showing cardiomyocyte hypertrophy on heart slices of AARS2^cKO^ group compared with AARS2^fl/fl^ group at 28 days after tamoxifen induction (scale bar, 100 μm; n = 5). (L) Masson staining and quantitative analysis showing increased cardiac fibrosis in AARS2^cKO^ hearts compared with AARS2^fl/fl^ control hearts at 28 days after tamoxifen induction (scale bar, 1 mm; n = 8). Mean ± s.e.m.

We then asked whether AARS2 cKO mice have abnormal heart function. By applying intraperitoneal injection of tamoxifen to induce Cre-LoxP mediated cardiac specific deletion of AARS2 for 5 days, we measured cardiac function by echocardiography (ECHO) at various time points and myocardial fibrosis by Masson staining at the experimental endpoint (Figure 1E). ECHO showed that ejection fraction (EF) and fractional shortening (FS) were comparable between AARS2^fl/fl^ control and AARS2^cKO^ mice before tamoxifen induction. At 7 days after tamoxifen induction, AARS2^cKO^ mice showed reduced EF and FS, with a downward trend on days 14 and 28 after induction, whereas AARS2^fl/fl^ control mice exhibited normal EF and FS (Figure 1G). Moreover, the heart wave tracings by ECHO were flatter in AARS2^cKO^ mice than those of AARS2^fl/fl^ mice at day 28 (Figure 1F). In addition, the AARS2^cKO^ mice performed worse in terms of cardiac output, as well as left ventricle end-diastolic diameter (LVEDD), and left ventricle end-systolic diameter (LVESD) (Figure 1H–I). At day 28 after induction, 50% of AARS2^cKO^ mice died, while all AARS2^fl/fl^ mice survived (Figure 1J). Wheat germ agglutinin (WGA) staining showed that cardiomyocyte hypertrophy was evident in AARS2^cKO^ hearts (Figure 1K), and Masson staining indicated severe cardiac fibrosis in AARS2^cKO^ hearts compared with AARS2^fl/fl^ hearts (Figure 1L) at 28 days after induction. The results collectively indicate that the deletion of AARS2 in cardiomyocytes results in abnormal cardiac function and fibrosis in mice, displaying characteristics of cardiomyopathy.

### AARS2 is required for energy metabolism in cardiomyocytes

In order to clarify the underlying mechanisms in AARS2^cKO^ hearts, we investigated cardiomyocyte apoptosis and metabolism. TUNEL staining showed that AARS2^cKO^ hearts had a significant increase of TUNEL^+^ cardiomyocytes compared with those in AARS2^fl/fl^ control hearts (Figure 2A). Consistently, the anti-apoptotic protein Bcl-2 decreased while the pro-apoptotic protein BAX increased in AARS2^cKO^ hearts (Figure 2B). Given that the depletion of AARS2 causes cellular energy metabolism imbalance in human patients (Zhou et al., 2019), we asked if AARS2^cKO^ mouse cardiomyocytes exhibit metabolic defects. We performed Seahorse assays to evaluate mitochondrial OXPHOS and cardiomyocyte glycolysis from adult mouse hearts, as well as the OXPHOS and glycolysis of NRCMs. We found that the baseline of mitochondrial oxygen consumption rate (OCR) in AARS2^cKO^ cardiomyocytes was similar to that in AARS2^fl/fl^ control cardiomyocytes, but the maximal OCR decreased in mutant mitochondria (Figure 2C). Furthermore, the baseline of extracellular acidification rate (ECAR) in AARS2^cKO^ cardiomyocytes was also similar to that of AARS2^fl/fl^ control cardiomyocytes, but the maximal ECAR decreased in mutant cardiomyocytes (Figure 2D). In addition, knocking down of AARS2 by small interfering RNA (siRNA) in NRCMs (Figure 2–figure supplement 1A–B) resulted in similar effect on both maximal OCR and ECAR *in vitro* (Figure 2E–F). Taken together, these results indicate that depletion of AARS2 in cardiomyocytes leads to cardiomyocyte death and impairs energy metabolism.

**Figure 2.**
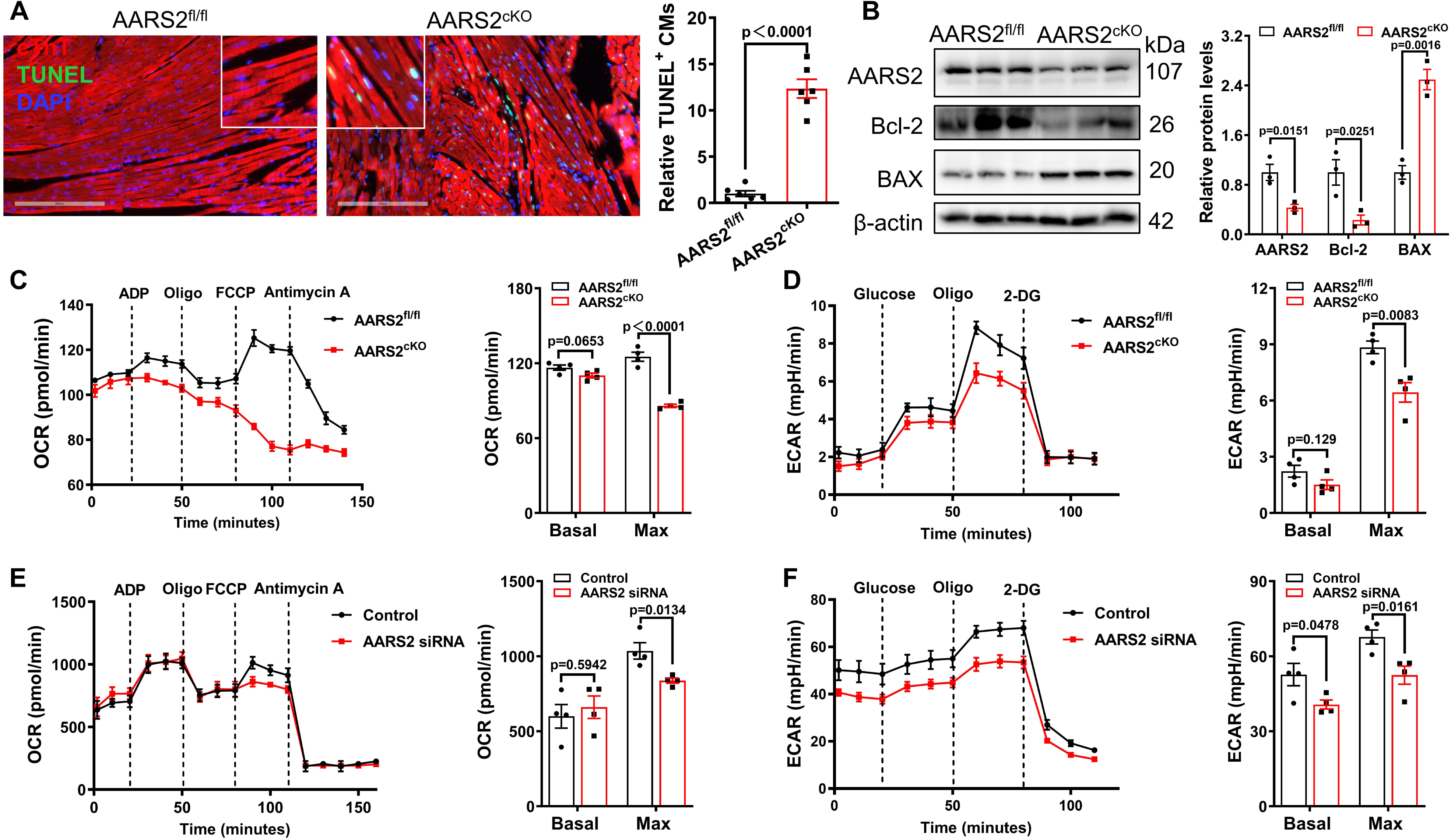
Cardiomyocyte-specific cKO of AARS2 results in cardiomyocyte apoptosis and energy metabolism deficiency. (A–B) Immunofluorescence staining showing increased numbers of cTnT^+^ TUNEL^+^ cardiomyocytes (A) and Western blot showing reduced anti-apoptotic protein Bcl-2 and increased pro-apoptotic protein BAX (B) in AARS2^cKO^ hearts compared with AARS2^fl/fl^ control hearts at 28 days post tamoxifen treatment (scale, 200 μm; n = 6 for panel A; n = 3 for panel B). (C) Seahorse analysis showing reduced oxygen consumption rate (OCR) of cardiac mitochondria in AARS2^cKO^ hearts compared with AARS2^fl/fl^ control hearts at 28 days after tamoxifen induction (n = 4). (D) Seahorse analysis showing reduced extracellular acidification rate (ECAR) of adult mouse cardiomyocytes in AARS2^cKO^ hearts compared with AARS2^fl/fl^ control hearts at 28 days after tamoxifen induction (n = 4). (E–F) Seahorse analysis showing reduced OCR (E) and ECAR (F) of NRCMs in AARS2 siRNA group compared with control group at 3 days after transfection (n = 4). Mean ± s.e.m.

### Overexpression of AARS2 in cardiomyocytes alleviates MI in mice

To further elucidate the function of AARS2 in the pathology of MI, we generated cardiomyocyte-specific overexpression of AARS2 driven by the α-MHC promoter by crossing α-MHC-MerCreMer mice with AARS2 transgenic mice (Figure 3A). In all experiments presented here, we used α-MHC-MerCreMer (AARS2^WT^) mice as control group while used α-MHC-MerCreMer; AARS2^Tg/+^ (AARS2^Tg/+^) mice as experimental group (Figure 3–figure supplement 1A). Western blots showed that AARS2 was only overexpressed in the heart after tamoxifen induction for 5 days while normally expressed in the liver, lung and skeletal muscle (Figure 3B and Figure 3–figure supplement 1C). In addition, immunofluorescence analysis revealed that AARS2 was highly expressed in cardiomyocytes of AARS2^Tg/+^ mouse hearts, whereas there was a significantly lower level of AARS2 in cardiomyocytes of AARS2^WT^ control hearts (Figure 3–figure supplement 1B). Together, we have successfully generated transgenic mice with cardiomyocyte-specific overexpression of AARS2.

**Figure 3.**
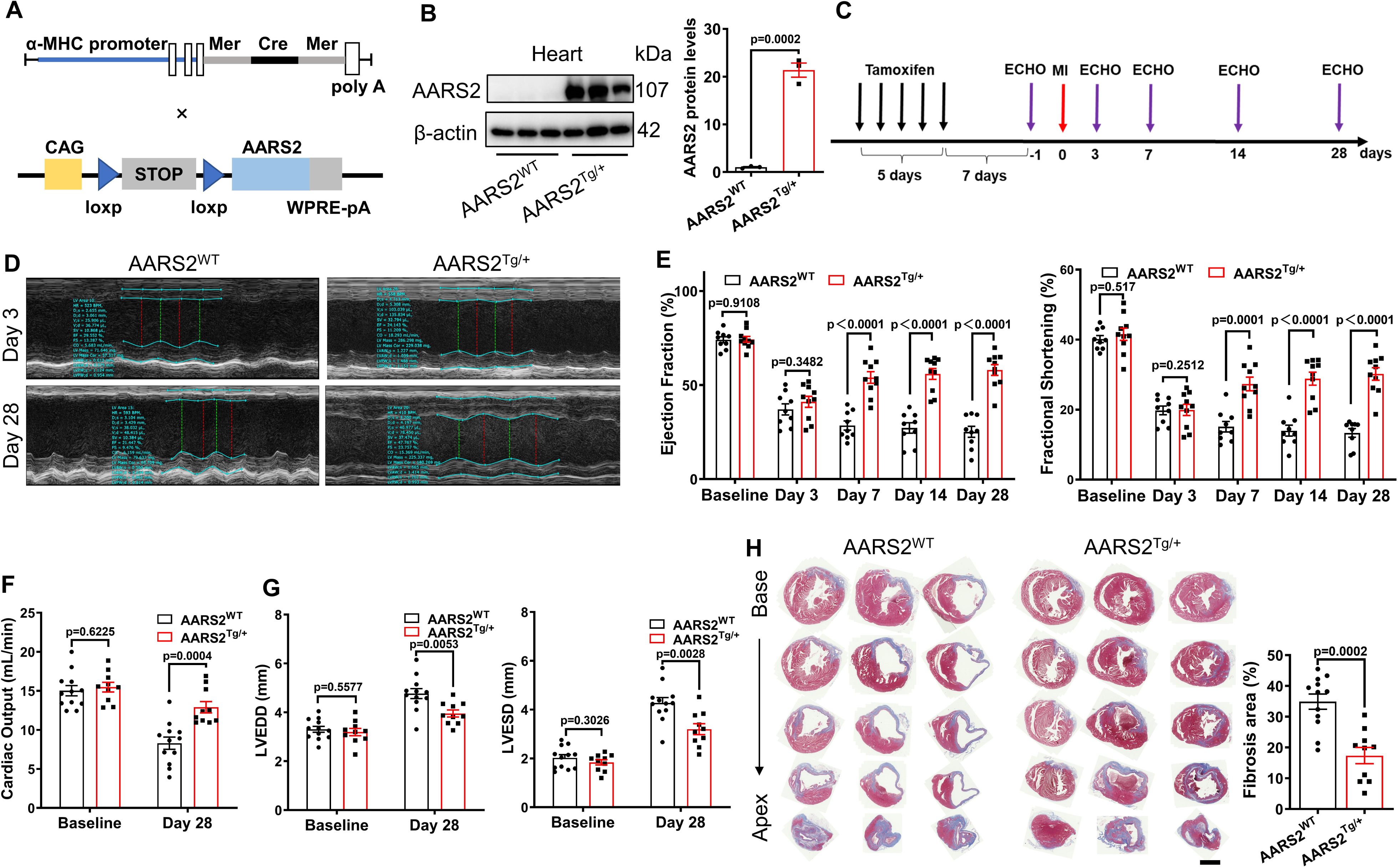
Cardiomyocyte-specific AARS2 overexpression improves cardiac function and decreases cardiac fibrosis in mice post-MI. (A) Schematic diagram of α-MHC-MerCreMer, and CAG-AARS2 mice that is driven by the CAG promoter. (B) Western blots showing transgenic overexpression of AARS2 proteins in the hearts of AARS2^Tg/+^ compared with AARS2^WT^ control mice (n = 3). (C) Experimental protocols for Tamoxifen induction for 5 days, and then recovery for 7 days before ECHO and MI. (D) Representative M-mode of ECHO in control and AARS2^Tg/+^ mouse hearts at 3 days or 28 days post-MI. (E) EF and FS of the AARS2^WT^ and AARS2^Tg/+^ mouse hearts were measured at different time points before and after MI (n = 10–11). (F) The cardiac output of AARS2^WT^ and AARS2^Tg/+^ mice was measured before MI and 28 days after MI (n = 10–11). (G) LVEDD and LVESD of AARS2^WT^ and AARS2^Tg/+^ mice before MI and 28 days after MI (n = 10–11). (H) Masson’s staining showing decreased fibrotic area in the hearts of AARS2^Tg/+^ compared with AARS2^WT^ mice at 28 days after MI (scale bar, 1 mm, n = 10–11). Mean ± s.e.m.

Then, we evaluated the impact of AARS2 overexpression on cardiac function post-MI by using AARS2 transgenic mice (Figure 3C). Before MI, we confirmed that EF and FS were comparable between AARS2^WT^ control and AARS2^Tg/+^ transgenic groups, and at day 3 after MI, EF and FS decreased significantly in both AARS2^WT^ control and AARS2^Tg/+^ transgenic mouse groups. ECHO showed that the waveforms of AARS2^Tg/+^ transgenic hearts recovered well compared with those of AARS2^WT^ control hearts at day 28 (Figure 3D). Importantly, from day 7 to 28 post-MI, we found that EF and FS gradually increased in AARS2^Tg/+^ transgenic group, while decreased in AARS2^WT^ control group (Figure 3E). In addition, the cardiac output increased, while LVEDD and LVESD decreased in AARS2^Tg/+^ transgenic hearts compared with AATS2^WT^ control hearts at day 28 post-MI (Figure 3F–G). Consistently, Masson staining showed that cardiac fibrosis decreased in the AARS2^Tg/+^ transgenic hearts compared with AARS2^WT^ control hearts at day 28 post-MI (Figure 3H). Briefly, these results suggest that overexpression of AARS2 in cardiomyocytes improves cardiac function and alleviates fibrosis in mice post-MI.

We next asked whether overexpression of AARS2 in cardiomyocytes has an impact on cardiomyocyte proliferation and coronary artery regeneration post-MI. We found that the proliferation index of either Ki67^+^/cTnT^+^ or pH3^+^/cTnT^+^ cardiomyocytes was comparable between AARS2^Tg/+^ transgenic and AARS2^WT^ control hearts at 7 days post-MI (Figure 3–figure supplement 2A–B). Similarly, overexpression of AARS2 in NRCMs had no effect on either Ki67^+^/cTnT^+^ or pH3^+^/cTnT^+^ cardiomyocytes *in vitro* (Figure 3–figure supplement 2C–D). As compensatory collateral vessel formation in the infarcted area contributes to cardiac repair, we evaluated the coronary vessel density using anti-CD31 (an endothelial cell marker) and α-SMA (a smooth muscle cell marker). However, there were no changes in coronary vessel density between AARS2^WT^ control and AARS2^Tg/+^ transgenic hearts at 7 days post-MI (Figure 3–figure supplement 2E). Additionally, WGA staining showed that AARS2 overexpression had no effect on cardiomyocyte hypertrophy in AARS2 transgenic hearts (Figure 3–figure supplement 2F). In conclusion, these results suggest that overexpression of AARS2 has no direct influence on cardiomyocyte proliferation, hypertrophy, and arterial vessel formation during the pathology of MI.

### Overexpression of AARS2 decreases cardiomyocyte mtROS and apoptosis

Given that overexpression of AARS2 does not promote cardiomyocyte proliferation and hypertrophy in adult mice, we further investigate whether it salvages cardiomyocytes from ultimate death. TUNEL staining revealed less amounts of TUNEL^+^/cTnT^+^ cardiomyocytes in AARS2^Tg/+^ transgenic hearts compared with those in AARS2^WT^ control hearts at 7 days post-MI (Figure 4A). Consistently, we found that the anti-apoptotic protein Bcl-2 increased, and pro-apoptotic protein BAX decreased in the left ventricles of AARS2^Tg/+^ transgenic hearts compared with AARS2^WT^ control hearts (Figure 4B). The level of lactate dehydrogenase (LDH) in the blood serum is a standard marker for assessing the severity of MI, therefore we examined LDH and found that it decreased in AARS2^Tg/+^ transgenic mouse serum (Figure 4C). In parallel, we also found that under hypoxia and reoxygenation (H/R), mitochondrial ROS indicator mtSOX decreased in AARS2 overexpression (AARS2^OE^) NRCMs compared with Mock NRCMs (Figure 4D). Consistent with the findings *in vivo*, AARS2^OE^ also decreased pro-apoptotic protein BAX while increased anti-apoptotic protein Bcl-2 in NRCMs subjected to H/R injury (Figure 4E), as well as decreased the release of LDH in the NRCM supernatants under either normoxia or hypoxia conditions (Figure 4F). Together, these results suggest that AARS2 overexpression alleviates ischemic- or hypoxia-induced cardiomyocyte damage.

**Figure 4.**
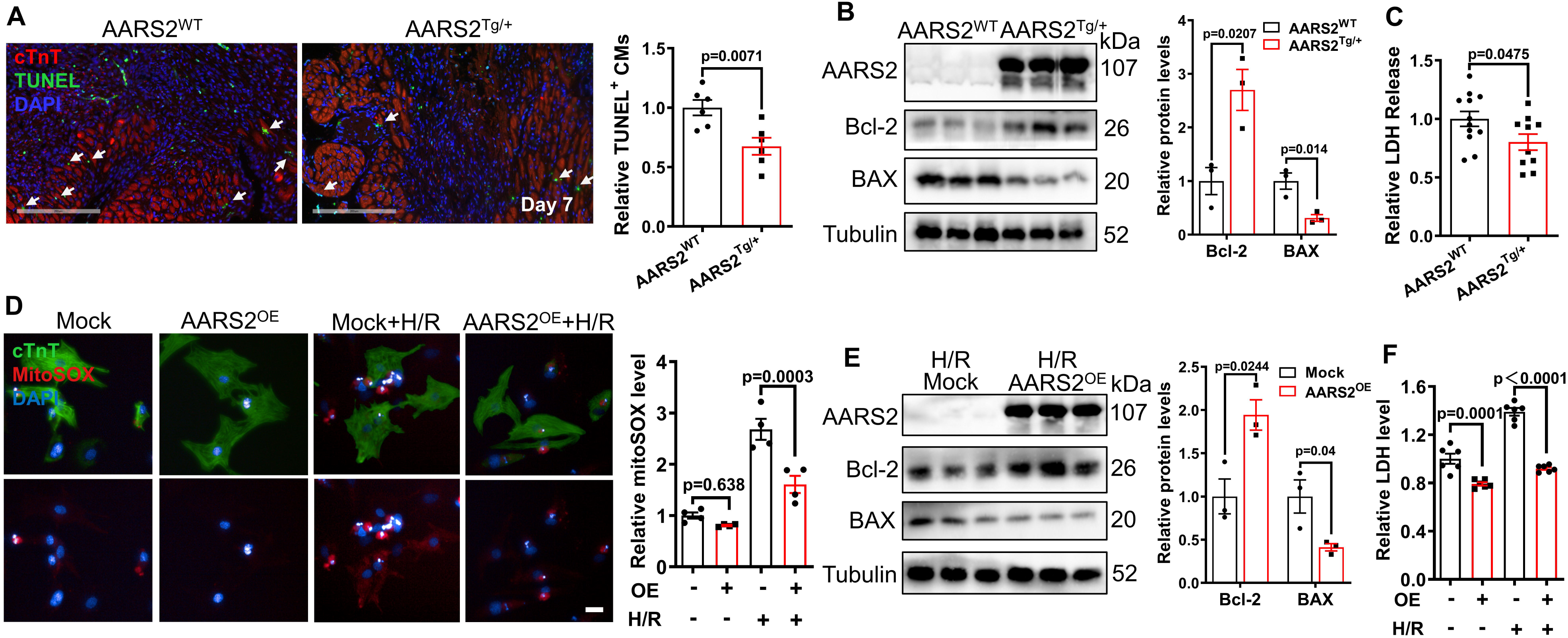
Overexpression of AARS2 attenuates cardiomyocyte apoptosis. (A) Immunofluorescence staining showing reduced cTnT^+^/TUNEL^+^ cardiomyocytes in AARS2^Tg/+^ hearts compared with AARS2^WT^ control hearts at 7 days after MI (scale bar, 200 μm, n = 6). (B) Western blots showing increased anti-apoptotic protein Bcl-2 and decreased pro-apoptotic protein BAX in AARS2^Tg/+^ compared with AARS2^WT^ control hearts at 7 days after MI (n = 3). (C) The serum level of LDH decreased in AARS2^Tg/+^ hearts compared with AARS2^WT^ control hearts at 28 days after MI (n = 10). (D) Immunofluorescence staining and quantitative analysis showing reduced MitoSOX in AARS2^OE^ NRCMs after 12 h of hypoxia followed by 1 h of reoxygenation (H/R, scale bar, 20 μm; n = 4). (E) Western blots showing increased Bcl-2 and decreased BAX in NRCMs overexpressing AARS2 (AARS2^OE^) compared with control NRCMs after 12 h of hypoxia followed by 1 h of reoxygenation (n = 3). (F) The level of LDH decreased in AARS2^OE^ NRCMs after 12 h of hypoxia followed by 1 h of reoxygenation (n = 6). Mean ± s.e.m.

### Overexpression of AARS2 in cardiomyocytes promotes glycolysis

We then asked what are the underlying mechanisms of AARS2 on cardiac protection. Since a previous study has shown that overexpression of AARS2 leads to intracellular metabolic changes (Zhao et al., 2018), we performed Mass Spectrometry for assessing metabolites of homogenates from heart tissues or NRCMs. We found that the amounts of lactate and pyruvate increased, while acetyl-CoA decreased in AARS2^Tg/+^ transgenic hearts compared with AARS2^WT^ control hearts (Figure 5A). Similarly, we also found that AARS2 overexpression led to elevating lactate and pyruvate as well as decreasing acetyl-CoA in AARS2^OE^ NRCMs compared with Mock control NRCMs (Figure 5B). To further assess the impact of AARS2 overexpression on cardiac energy metabolism, we conducted Seahorse to evaluate mitochondrial OXPHOS and cellular glycolysis in adult mouse hearts and NRCMs. The baseline of OCR was comparable between AARS2^Tg/+^ transgenic and AARS2^WT^ control hearts, but the maximal OCR decreased in AARS2^Tg/+^ transgenic hearts (Figure 5C). Additionally, the baseline of ECAR had no difference in adult cardiomyocytes of both groups, but the maximal ECAR increased in AARS2^Tg/+^ transgenic cardiomyocytes compared with AARS2^WT^ control cardiomyocytes (Figure 5D). We found similar findings that the baseline OCR and ECAR had no changes in both groups, but the maximal OCR decreased and maximal ECAR increased in AARS2^OE^ NRCMs (Figure 5E–F). Thus, these results indicate that AARS2^OE^ in cardiomyocytes can shift the metabolic profile from OXPHOS to glycolysis. Under ischemic or hypoxia conditions, this metabolic shift allows cardiomyocytes to avoid OXPHOS dysfunction and enables them to produce energy through glycolysis as a protective mechanism for self-preservation and energy supply. This metabolic adaptation may be crucial in maintaining cell viability and function during the periods of reduced oxygen availability in the heart.

**Figure 5.**
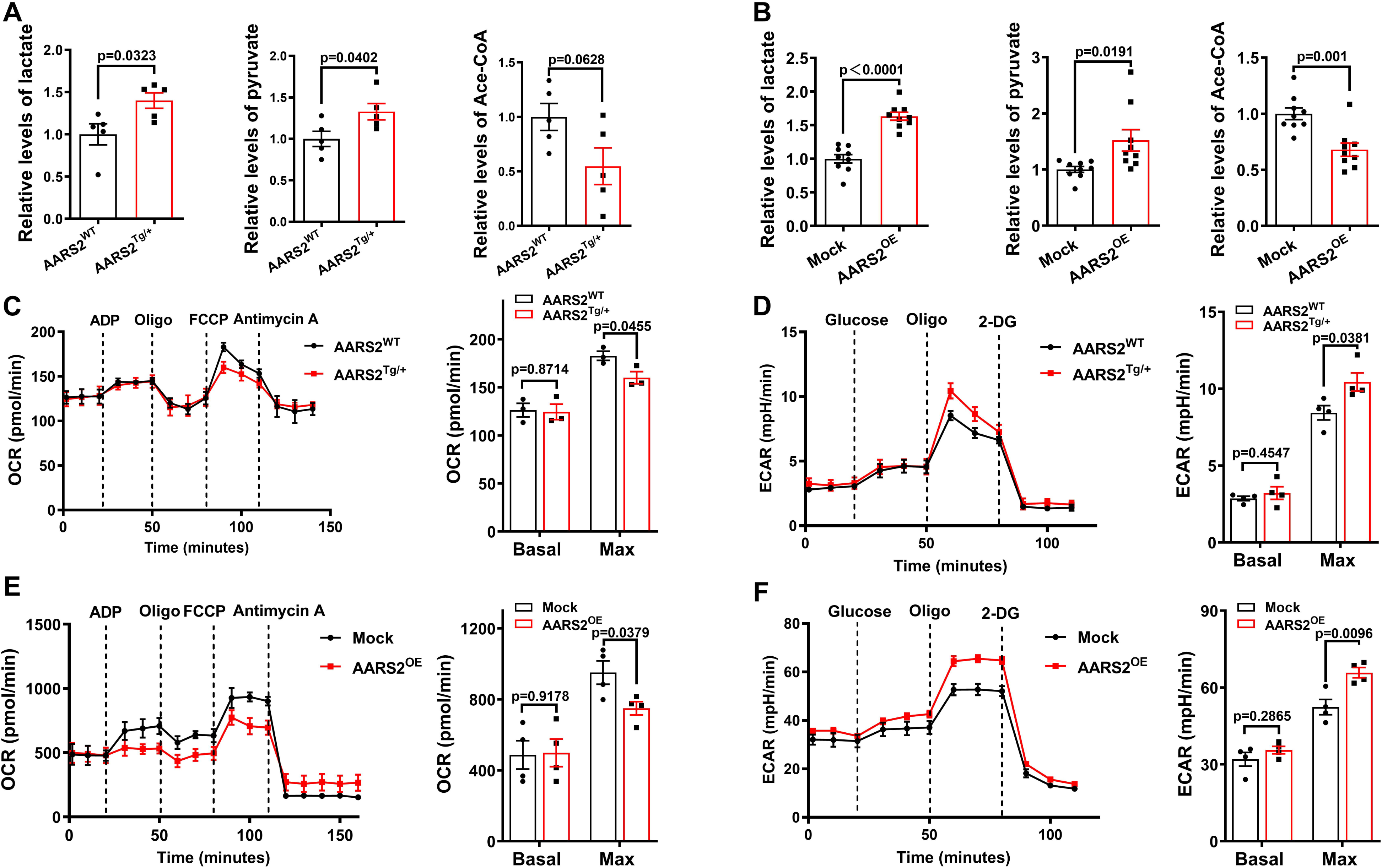
Cardiomyocyte overexpression of AARS2 regulates cardiac metabolism. (A) Mass spectrometry showing increased lactate and pyruvate but reduced acetyl-CoA in AARS2^Tg/+^ hearts compared with AARS2^WT^ hearts after 7 days of MI (n = 5). (B) Mass spectrometry showing increased lactate and pyruvate but decreased acetyl-CoA in NRCMs overexpressing AARS2 (AARS2^OE^) for 3 days (n = 9). (C) Seahorse analysis showing OCR of cardiac mitochondria in AARS2^WT^ and AARS2^Tg/+^ mice at 28 days after tamoxifen induction (n = 3). (D) Seahorse analysis showing ECAR and quantitative analysis of adult mouse cardiomyocytes in AARS2^WT^ and AARS2^Tg/+^ mice at 28 days after tamoxifen induction (n = 4). (E–F) Seahorse analysis showing OCR (E) and ECAR (F) of NRCMs in Mock control and AARS2^OE^ groups at 3 days after transfection (n = 4). Mean ± s.e.m.

### AARS2 promotes glycolysis *via* regulating the PKM2 translation and ratio of PKM2 dimers to tetramers

We then asked how elevated AARS2 in cardiomyocytes leads to metabolic changes from OXPHOS toward glycolysis. AARS2 is an alanyl-tRNA synthase that is critical for RNA translation, so we applied Ribosome RNA-seq to examine whether over-expression of AARS2 has any effect on RNA translation of metabolic genes. We found that AARS2 overexpression promoted the translation of glycolytic proteins, including PKM (Figure 6A), PDK4 and LDHA proteins (Figure 6B). Considering that PKM is the key enzyme that promotes glycolysis (Tang et al., 2023; Vander Heiden, Cantley, & Thompson, 2009), which PKM1 and PKM2 are two different splicing forms of the PKM gene (Dayton, Jacks, & Vander Heiden, 2016), we examined whether AARS2 overexpression has effect on PKM1 and PKM2 translation. Western blots showed a significant increase in PKM2 proteins with AARS2 overexpression in both mouse hearts and NRCMs (Figure 6D–E). Conversely, PKM2 protein markedly decreased in AARS2^cKO^ hearts (Figure 6F). On the other hand, we found that AARS2 overexpression had no effect on PKM1 proteins (Figure 6C), suggesting that AARS2 overexpression upregulates PKM2 but not PKM1 translation. In addition, overexpression of AARS2 also enhanced the translation of other signaling pathways response to hypoxia, and sodium ion transport (Figure 6–figure supplement 1A–D), which has a positive effect on mitochondrial function and cardiomyocyte function.

**Figure 6.**
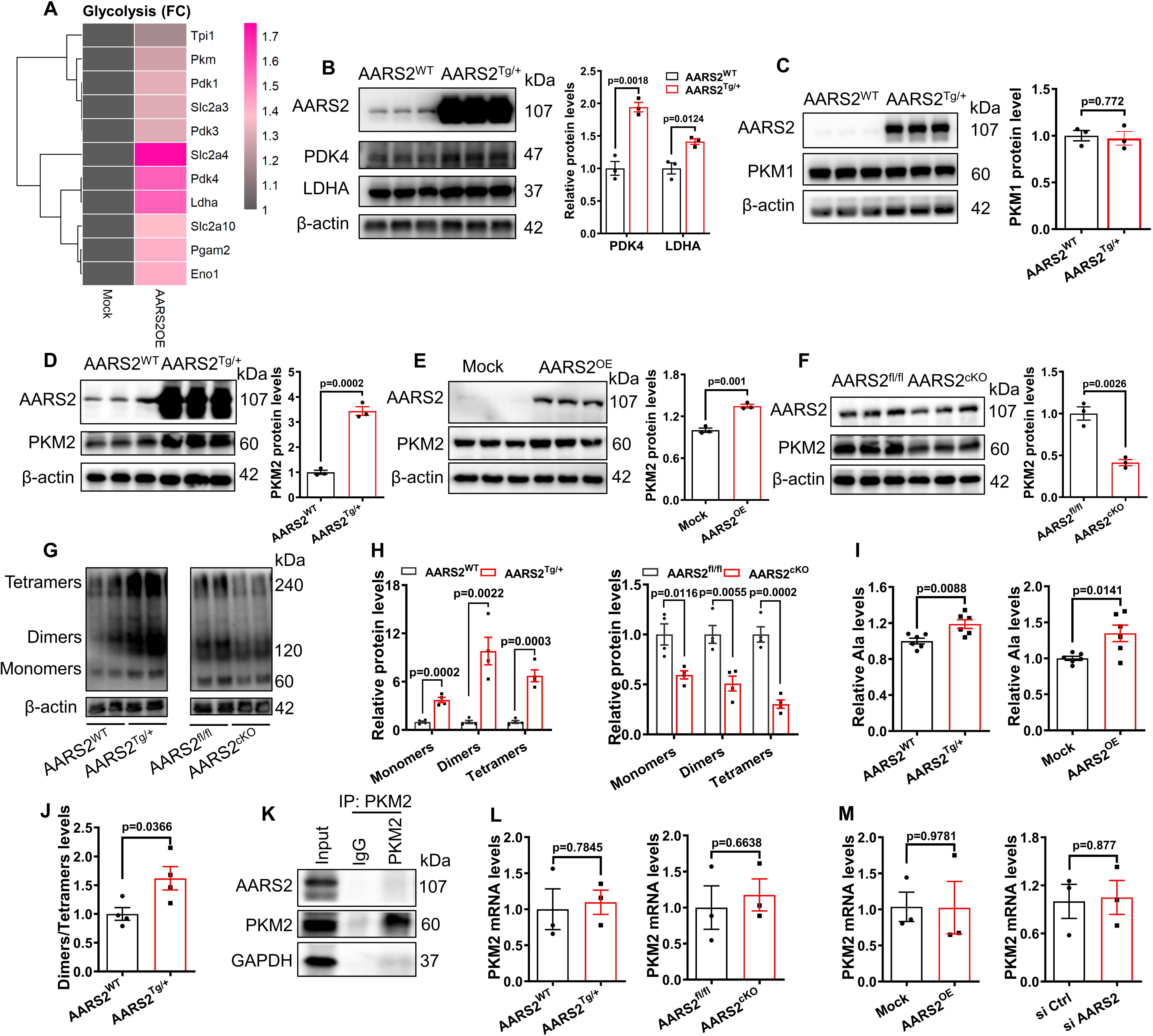
Overexpression of AARS2 increases the protein level of glycolytic PKM2 *via* enhancing PKM2 translation. (A) Ribosome RNA-Seq showing elevated translation of signaling pathways of glycolysis in the AARS2^OE^ NRCMs compared to the Mock NRCMs. (B) Western Blots showing the level of AARS2, PDK4 and LDHA proteins in the hearts of AARS2^WT^ control and AARS2^Tg/+^ transgenic mice (n = 3). (C) Western Blots showing the level of AARS2 and PKM1 proteins in the hearts of AARS2^WT^ control and AARS2^Tg/+^ transgenic mice (n = 3). (D–F) Western Blots showing the level of AARS2 and PKM2 proteins in the hearts of AARS2^WT^ control and AARS2^Tg/+^ transgenic mice (n = 3) (D), in Mock control and AARS2^OE^ NRCMs (E) (n = 3), and in the hearts of AARS2^fl/fl^ and AARS2^cKO^ mice (F) (n = 3). (G–H) Western Blots by non-denatured gels (G) and statistics (H) showing the amounts of PKM2 monomers, dimers and tetramers in the hearts of AARS2^WT^ control and AARS2^Tg/+^ transgenic mice, and in the hearts of AARS2^fl/fl^ and AARS2^cKO^ mice (n = 4). (I) Mass spectrometry analysis measuring the amounts of alanine (Ala) from homogenates of heart tissues (n = 6) and NRCM lysates (n = 6). (J) Ratio of quantitative results of PKM2 dimers and tetramers in the hearts of AARS2^WT^ control and AARS2^Tg/+^ transgenic mice of panel H (n = 4). (K) Co-immunoprecipitation reveals no evident interactions between PKM2 and AARS2 in NRCMs. (L–M) qRT-PCR showing the comparative level of PKM2 mRNA in the hearts of control sibling and AARS2^Tg/+^ transgenic hearts; control sibling and AARS2^cKO^ hearts (L); and in control, AARS2^OE^, and AARS2^siRNA^ NRCMs (M) (n = 3). FC, Fold changes; Mean ± s.e.m.

Because PKM2 can be converted between dimers and tetramers, which is critical for maintaining energy metabolism between glycolysis and OXPHOS, with dimers toward glycolysis (Dayton, Jacks, et al., 2016). We found that AARS2 overexpression increased, while AARS2 cKO decreased, PKM2 dimers and tetramers (Figure 6G–H), thus corroborating the increase of glycolysis in AARS2 transgenic hearts and the reduction of glycolysis and OXPHOS in AARS2 cKO. Previous studies have suggested that an increase in alanine causes a shift of PKM2 tetramers to dimers, consequently promoting glycolysis (Dayton, Jacks, et al., 2016). Mass spectrometry analysis revealed that alanine indeed increased in transgenic hearts and NRCMs upon AARS2 overexpression (Figure 6I), which is consistent with the observation that the ratio of dimers/tetramers increased in AARS2 transgenic hearts (Figure 6J). In addition, co-immunoprecipitation assay revealed no direct interaction between AARS2 and PKM2 (Figure 6K). Moreover, regardless of AARS2 overexpression or cKO in mouse hearts and NRCMs, we found that the level of PKM2 mRNA was not altered (Figure 6L–M), suggesting that AARS2 has no effect on PKM2 transcription. Therefore, AARS2 overexpression enhances glycolysis through regulating alanine and PKM2 translation.

### Activation of PKM2 alleviates cardiomyopathy in AARS2 cKO mice

To establish the functional relationship between AARS2 and PKM2, we then investigated whether the PKM2 activator TEPP-46 (TEPP-46 tetramerizes PKM2, thereby enhancing energy supply) alleviates cardiomyocyte death caused by AARS2 deficiency. After inducing AARS2 deficiency with tamoxifen in AARS2^cKO^ mice, we administered TEPP-46 according to the timeline and examined cardiac function by ECHO as shown (Figure 7A). We found that the baseline of EF and FS were comparable before tamoxifen induction, EF and FS decreased at day 0 (5 days after tamoxifen induction) in both vehicle (AARS2^cKO^) control and TEPP-46 (AARS2^cKO^) groups. While EF and FS continued to decline in vehicle control mice, EF and FS exhibited a significant recovery and tended to stabilize in TEPP-46 mice after 7 days of treatment (Figure 7B). ECHO revealed better waveforms and an improvement in cardiac function in TEPP-46 group compared to vehicle group at 28 days post treatment (Figure 7C). Moreover, increased cardiac output, decreased LVESD and LVEDD indicated a significant improvement in TEPP-46 group (Figure 7D–E). Masson staining showed less cardiac fibrosis in the hearts of TEPP-46 group compared with vehicle group at 28 days post treatment (Figure 7F). Furthermore, AARS2 knockdown in NRCMs resulted in a significant increase in LDH *in vitro*, which was repressed by TEPP-46 treatment (Figure 7G). Additionally, TEPP-46 also effectively suppressed mitochondrial ROS generation induced by AARS2 knockdown (Figure 7H). Taken together, these results suggested that activating PKM2 by

**Figure 7.**
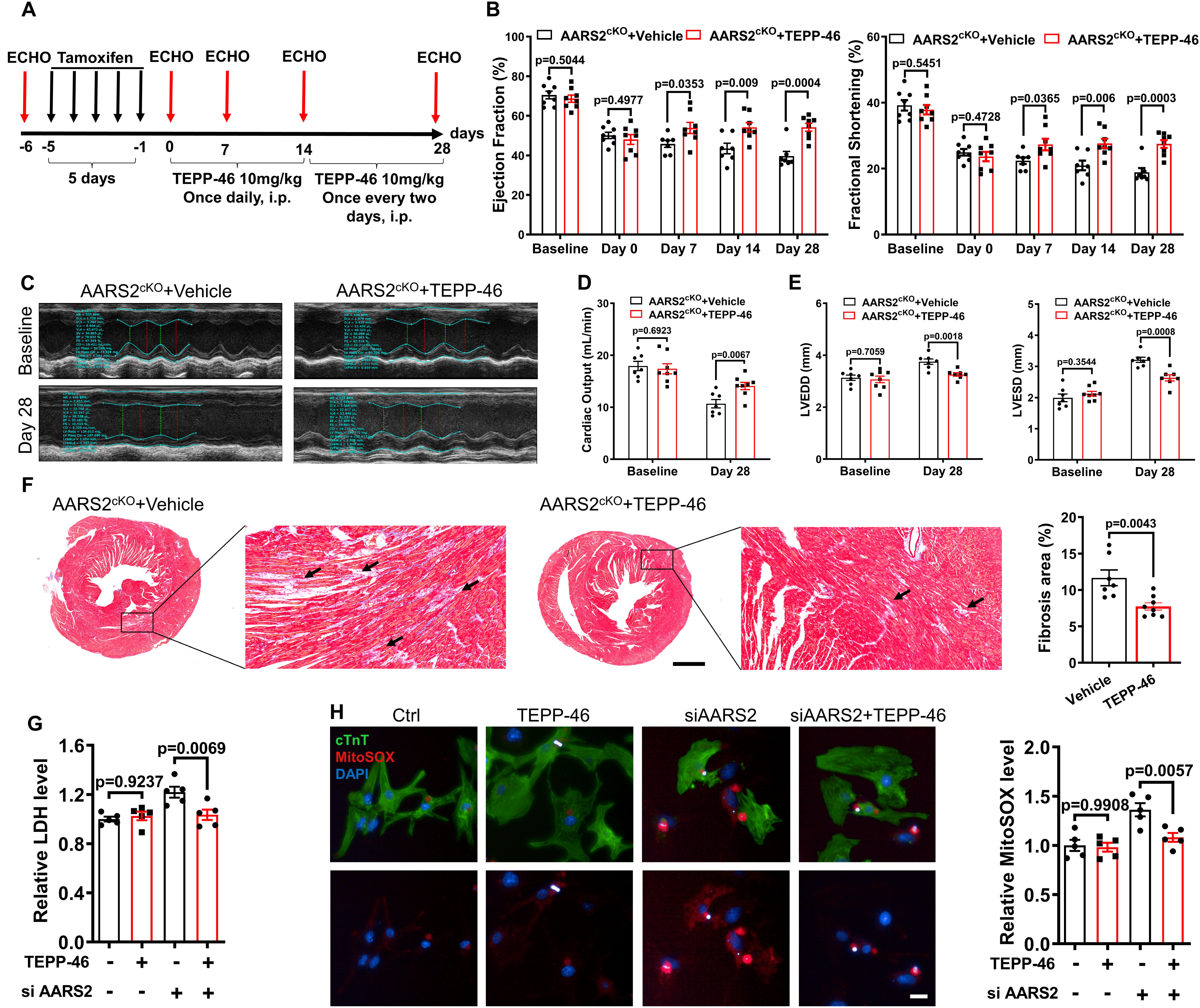
PKM2 activator TEPP-46 improves cardiomyopathy in AARS2 cKO mice. (A) Experimental scheme and time points for tamoxifen induction, ECHO and TEPP-46 administration. (B) EF and FS of AARS2^cKO^ mouse hearts at different time points after administration of control solvent and TEPP-46 (n = 7–8). (C) Representative M-mode of ECHO in different groups of 4-week mice. (D) Cardiac outputs of AARS2^cKO^ mouse hearts at different time points after administration of control solvent and TEPP-46 (n = 7–8). (E) Left ventricular end-diastolic diameter (LVEDD) and left ventricular end-systolic diameter (LVESD) at 28 days of AARS2^cKO^ mice after administration of control solvent and TEPP-46 (n = 7–8). (F) Masson staining showing cardiac fibrosis of AARS2^cKO^ mouse hearts at 28 days after administration of either control solvent or TEPP-46 (scale bar, 1 mm; n = 7–8). (G) Measurements of LDH release in NRCMs from the control group and TEPP-46 group (20 μM) after AARS2 siRNA or control siRNA treatment for 72 h (n = 5). (H) Quantitative analysis of MitoSOX immunofluorescence in NRCMs from the control group and TEPP-46 group (20 μM) after AARS2 siRNA or control siRNA treatment for 72 h (scale bar, 20 μm; n = 5). Mean ± s.e.m.

## Discussion

CVDs have become a prevalent cause of morbidity and mortality worldwide (Z. Zhang et al., 2023). The pathological processes of CVDs involve irreversible cardiomyocyte death, oxidative stress, inflammation, and cardiac fibrosis, which further contribute to adverse cardiac remodeling and ultimately lead to heart failure. Currently, there are no effective therapeutic strategies available for CVDs. This study showed that AARS2 plays a crucial role in salvaging cardiomyocytes from ischemic stress *via* fine-tuning PKM2-mediated energy metabolism, thus presenting novel therapeutic targets and potential small molecules for treating cardiomyopathy and MI.

Previous studies have shown that AARS2 mutations lead to abnormal cardiomyocyte function, eventually to cardiomyopathy in human patients (Nielsen et al., 2020). However, it remains unknown whether AARS2 functions in cardiomyocytes or non-cardiomyocytes and how AARS2 regulates cardiomyocyte function and metabolism. Mitochondrial OXPHOS is the primary pathway for adult cardiomyocytes to produce ATP, relying on oxygen in the mitochondria to oxidize substrates and generate ATP through the cellular respiratory chain. Under hypoxia, glycolysis serves as an alternative pathway for cells to produce energy, breaking down glucose into lactate and generating a small amount of ATP (Vander Heiden et al., 2009; Wei et al., 2023). By creating cardiomyocyte-specific cKO and transgenic overexpression of AARS2 mice, we found that AARS2 is not only required for cardiomyocyte function, but also improving cardiomyocytes from ischemia and hypoxia, thus establishing the critical function of AARS2 in cardiomyocytes. This provides a principle for future AARS2 gene therapy with targeted delivery into cardiomyocytes not non-cardiomyocytes. Our mechanistic investigations revealed that mitochondrial OXPHOS and glycolysis were impaired in cardiomyocytes lacking AARS2, leading to energy supply disruption and metabolic imbalance, ultimately resulting in cardiomyocyte death. On the other hand, AARS2 overexpression had a striking protective effect on cardiomyocytes, partly *via* enhancing glycolysis and resistance of cardiomyocytes to ischemia and hypoxia. Unbiased Ribosome RNA-Seq and functional analyses point out PKM2 as an important mediator of AARS2 in regulating cardiomyocyte function and metabolism. PKM2 decreased in the hearts of AARS2 cKO mice, with the reduced formation of PKM2 dimers and tetramers; and AARS2 overexpression led to an increase in glycolysis and attenuated OXPHOS, with increased PKM2 translation and the ratio of PKM2 dimers to tetramers. PKM2 is a key enzyme in the glycolytic pathway, involved in converting PEP into pyruvate and ATP (Méndez-Lucas et al., 2017). Under certain conditions, an elevation in PKM2 levels can enhance the activity of the glycolytic pathway, thereby increasing the cell ability to produce ATP through glycolysis (Wong et al., 2015). Recent studies have found that not only AARS1 (Ju et al., 2024; Zong et al., 2024), but also AARS2 is a lactate binding protein (Mao et al., 2024). Increased AARS2 further leads to increased binding to lactate, and PKM2 dimer, as a key enzyme in lactate production, can promote the production of lactate through glycolysis. Thus, AARS2 overexpression promotes the glycolytic pathway by increasing PKM2 translation and formation of dimers, thereby supporting the survival and function of cardiomyocytes. This metabolic adaptation is crucial for the heart to maintain its physiological functions under hypoxic or ischemic stress. Indeed, the PKM2 activator TEPP-46 enables to rescue cardiomyopathy phenotypes including cardiomyocyte death and cardiac fibrosis. This is because TEPP-46 enhances the tetramerization of PKM2, which enhances OXPHOS for cardiomyocytes. Therefore, this work presents a new cardiomyocyte survival mechanism involving AARS2-PKM2 signaling under ischemic conditions.

It is widely recognized that cardiomyocytes undergo dramatic metabolic alterations and responses to oxidative stress during the pathogenesis of CVDs. Our Seahorse data suggested that the maximal OCR decreased, while the maximal ECAR strikingly increased, in AARS2^Tg/+^ transgenic hearts. These findings suggest an enhanced glycolytic pathway in AARS2-overexpressing cardiomyocytes, thereby augmenting their capacity for ATP production through glycolysis. This metabolic adaptation may contribute to sustaining the survival and functionality of cardiomyocytes, particularly under hypoxia or other stress conditions. This notion was also supported by functional analyses of AARS2 in NRCMs. Additionally, overexpression of AARS2 inhibits mtROS production and confers protection against ischemia and H/R-induced injury in cardiomyocyte damage and disease occurrence (Wei et al., 2023). Despite causing a reduction in OXPHOS, AARS2 overexpression also suppresses mtROS generation caused by ETC disorders, thus mitigating the detrimental effects of oxidative stress. Consequently, overexpression of AARS2 provides a protective mechanism by reducing mtROS levels and helps maintain normal physiological homeostasis within cardiomyocytes. In summary, upregulation or activation of PKM2 dimers mediated by AARS2 overexpression enhances glycolysis, thereby supporting the survival and functionality of cardiomyocytes, especially under hypoxic or stressful conditions. This metabolic adaptation coupled with reduction of ROS holds significant importance for maintaining normal cardiac function during challenging situations.

However, this study has certain limitations. The interaction between PKM2 and HIF-1α in the nucleus can affect the expression of glycolysis genes and promote cell survival (Chen et al., 2014; Palsson-McDermott et al., 2015; Prakasam, Iqbal, Bamezai, & Mazurek, 2018), thus it remains be addressed whether AARS2 acts through this mechanism in cardiomyocytes. Investigating specific interactions between PKM2 and AARS2 in the context of cardiac pathophysiology will provide a better understanding of cardiomyocyte metabolism and adaptation mechanisms, but glycolysis is only one of the processes that the body makes adaptive changes under special conditions. It remains unclear whether continuously excessive high-level of glycolysis is beneficial to cardiomyocytes. This study primarily relies on mouse models and NRCMs, lacking sufficient human data to validate the experimental outcomes in human cardiomyocytes. Therefore, more researches related to human subjects are necessary before translating these findings to clinical treatments in humans. Moreover, ribosome profiling sequencing (Ribo-Seq) data show that AARS2 enhances PKM2 translation only as part of the investigated mechanisms. In fact, AARS2 also affects the translation of many other proteins. Further investigations are needed to address the influence of AARS2 on gene translation. Additionally, delving into more mutual interactions and regulatory mechanisms between AARS2 and PKM2 will help uncover the regulatory mechanisms of AARS2 on cardiomyocyte energy metabolism, providing a deeper understanding and potential therapeutic targets for the treatment of CVDs. Therefore, future investigations are needed to address how AARS2 regulate mitochondrial function in cardiomyocytes, how AARS2 interacts with PKM2 complex to regulate mitochondrial metabolisms, and how the AARS2-PKM2 signaling is related to the pathogenesis of human cardiomyopathy.

## Materials and methods

| **Key Resources Table** |  |  |  |  |
| --- | --- | --- | --- | --- |
| **Reagent type (species) or resource** | **Designation** | **Source or reference** | **Identifiers** | **Additional information** |
| genetic reagent (M. musculus) | AARS2 flox/flox | Zhao Lab, Fudan University | N/A | C57BL/6J |
| genetic reagent (M. musculus) | AARS2 Transgenic | Zhao Lab, Fudan University | N/A | C57BL/6J |
| genetic reagent (M. musculus) | α-MHC-Mer CreMer | Xiong Lab, Peking University | <u>N/A</u> | C57BL/6J |
| primary cell ( Rat ) | Primary neonatal rat cardiomyocytes | Charles River | N/A | CD®(SD) IGS Rat, 3 days |
| antibody | anti-PKM2 (Rabbit polyclonal) | CST | 4053S | WB (1:1000) |
| antibody | anti-AARS2 (Rabbit polyclonal) | Abcam | ab197367 | WB (1:1000) |
| antibody | anti-Bcl2 (Rabbit polyclonal) | CST | 3498S | WB (1:1000) |
| antibody | anti-BAX (Rabbit polyclonal) | CST | 14796S | WB (1:1000) |
| antibody | anti-PKM1 (Rabbit polyclonal) | Proteintech | 15821-1-A P | WB (1:1000) |
| antibody | anti-PDK4 (Rabbit polyclonal) | Bioss | bs-0682R | WB (1:1000) |
| antibody | anti-LDHA<br>(Rabbit polyclonal) | Bioss | bs-34202R | WB (1:1000) |
| antibody | Goat Anti-Rabbit IgG (H&L)-HRP Conjugated | EASYBIO | BE0101-100 | WB (1:10000) |
| antibody | HRP-conjugated GAPDH | Proteintech | HRP-60004 | WB (1:10000) |
| antibody | HRP-conjugated Beta Actin | Proteintech | HRP-60008 | WB (1:10000) |
| antibody | HRP-conjugated Alpha Tubulin | Proteintech | HRP-66031 | WB (1:10000) |
| antibody | anti-Ki67<br>(Rabbit polyclonal) | CST | 12075S | IF (1:300) |
| antibody | anti-pH3<br>(Rabbit polyclonal) | CST | 53348S | IF (1:300) |
| antibody | anti-cTnT<br>(Mouse polyclonal) | Abcam | ab8295 | IF (1:300) |
| antibody | anti-α-SMA<br>(Mouse polyclonal) | CST | 48938S | IF (1:300) |
| antibody | anti-CD31<br>(Rabbit polyclonal) | Abcam | ab18291 | IF (1:300) |
| antibody | Goat Anti-Rabbit IgG H&L (Alexa Fluor® 488) | Abcam | ab150077 | IF (1:500) |
| antibody | Goat Anti-Mouse IgG H&L (Alexa Fluor® 555) | Abcam | ab150118 | IF (1:500) |
| recombinant DNA reagent | pLenti-CMV-MCS-PGK-Puro-WPRE (plasmid) | This paper | N/A | CMV-MCS-PGK-Puro |
| Ribo-seq data | Ribosome sequencing of the effect of control lentivirus and AARS2 overexpression on lentivirus on neonatal rat cardiomyocyte translation | ArrayExpress | E-MTAB-13767 | N/A |

### Experimental animals

Sprague-Dawley neonatal rats (postnatal day 3, P3) and adult mice were obtained from Vital River Laboratory Animal Technology Co., Ltd (Beijing, China). All experimental procedures involving animal subjects were conducted in accordance with the protocols approved by the Institutional Animal Care and Use Committee at Peking University, Beijing, China (IACUC: IMM-XingJW-4).

Generation of cardiomyocyte-specific AARS2 knockout mice and AARS2 overexpression mice were achieved by crossing either AARS2^fl/fl^ or AARS2^Tg^ mice with α-MHC-MerCreMer transgenic mice (Sohal et al., 2001). The AARS2^fl/fl^ mice and AARS2^Tg^ mice were obtained from Dr. Shi-Min Zhao’s lab at Fudan University, which were kept in C57BL/6J background.

The AARS2 cKO mice were given with TEPP-46 (10 mg/kg, MCE, Shanghai, China) by intraperitoneal injection once a day for the first and second weeks, and every other day for the third and fourth weeks after tamoxifen induction (20 mg/kg, Sigma-Aldrich, St. Louis, Missouri, USA).

### Adult mouse MI

The MI model was established in mice at an age ranging 8–10 weeks through ligation of the left anterior descending coronary artery (Du et al., 2022). Initially, each mouse was anesthetized by intraperitoneal injection of tribromoethanol (300 mg/kg; Sigma-Aldrich). Following complete anesthesia, the mouse was positioned supine and immobilized. Endotracheal intubation was performed, and the mouse was connected to a small-animal ventilator (MouseVent, Kent Scientific Corp., Torrington, CT, USA). The thoracotomy was carried out in the left intercostal space between the third and fourth ribs, exposing the heart by removing the pericardium. The left anterior descending coronary artery was permanently ligated using a 6-0 non-absorbable surgical suture, and the chest and skin were promptly sutured. Finally, the mouse was removed from the ventilator and kept warm until fully awaken.

### Echocardiography

ECHO was performed on anesthetized mice using a Vevo 3100 system (Visual Sonics, Toronto, ON, Canada) with 1.0% isoflurane anesthesia (T. Zhang et al., 2016). The hair on the left chest was completely removed using a depilatory paste to ensure optimal image acquisition. Two-dimensional ECHO images were obtained using a 20-MHz variable-frequency transducer in both the mid-ventricular short axis and the parasternal long axis. Heart rate, LVEDD and LVESD were measured from the short-axis M-mode tracings at the level of the papillary muscle. Left ventricle functional parameters, including the percentage of EF, FS, and cardiac output were calculated using the aforementioned measurements and accompanying software.

ECHO data were collected and analyzed on day -1 (baseline, 1 day before MI) as well as on days 3, 7, 14, and 28 post-MI.

### Pathological evaluation of mouse hearts

Mouse hearts were harvested and fixed in 4% paraformaldehyde (PFA) for a minimum of 2 days. Following fixation, the hearts underwent a series of processing steps, including dehydration, clearing, and infiltration, using a Histoprocessor (Tissue-Tek, Sakura, Tokyo, Japan). The paraffin-embedded tissues were then sectioned at a thickness of 5 µm. Masson’s trichrome staining was performed according to the protocol (Chuang et al., 2016). Briefly, heart sections were immersed in Bouin’s solution, followed by sequential staining with Mayer’s hematoxylin solution, Biebrich scarlet–acid fuchsin, phosphomolybdic acid–phosphotungstic acid, and aniline blue reagents (Sigma-Aldrich), with distilled water rinses between each step. Subsequently, the sections were dried, mounted on glass slides, and examined and photographed under a microscope.

### Isolation of NRCMs and transfection

NRCMs were isolated from P3 Sprague-Dawley rat hearts according to a previously described method (Du et al., 2022). Briefly, the hearts of rat pups were dissected and washed with Ca^2+^- and Mg^2+^-free HBSS (MacGene, Beijing, China). Using micro-dissecting scissors, ventricular tissues were minced into small pieces, and were then treated with 5 mL of digestion solution containing collagenase II (0.3 mg/mL; Thermo Fisher Scientific, Pittsburgh, PA, USA) and trypsin (1 mg/mL; Amresco, Pennsylvania, PA, USA) in HBSS for 5 min at 37°C. The cell supernatant was collected and the residual tissue was repeatedly treated with the digestion solution until little remained. The supernatants were transferred to a tube with an equal volume of ice-cold Dulbecco’s Modified Eagle Medium (DMEM; MacGene) containing 10% fetal bovine serum (FBS; Thermo Fisher Scientific, 16140071) and 1% penicillin-streptomycin, and then centrifuged at 400 g for 5 min. The cell pellets were re-suspended in 25-mL DMEM containing 10% FBS, 1% penicillin-streptomycin and 1 μmol/L cytosine arabinoside. The cells were incubated in a 100-mm dish for 1.5 h at 37°C to eliminate fibroblast contamination, then non-adherent cells were collected and seeded at a concentration of 5×10^5^ cells/mL. After incubation for 48-72 h, the medium was removed and the NRCMs were then cultured with DMEM containing 10% FBS and 1% penicillin-streptomycin for further analysis.

Transfection of cells was performed by using Lipofectamine 3000 (Invitrogen) according to the manufacturer’s protocol. Cells were harvested at 72 h post transfection and washed in phosphate-buffered saline for further RNA extraction and whole-cell protein extraction. The sequences of the small interfering RNA used are listed in Table S1.

The vector pLenti-CMV-MCS-PGK-Puro-WPRE was selected for lentiviral packaging, and the AARS2 gene was used according to the Gene Bank. Briefly, AARS2 sequence was ligated with restriction enzymes. The vector was ligated, and the sequence-confirmed plasmids were transfected into 293T cells. The supernatant was collected, and lentiviruses at 10 MOI were used to infect the cells for AARS2 overexpression. AARS2 overexpression and its control, Ad-green fluorescent protein lentiviruses, were utilized. NRCMs were infected with the lentivirus for 72 h. Then NRCMs were fixed for staining or lysed for protein extraction at 72 h after removing virus from NRCMs.

### Genotyping

The last 1–3 mm of the mouse tails was used for genotyping. Lyophilized proteinase K (1245680100, Merck) was dissolved at 10 mg/mL in 50 mmol/L Tris-HCl, pH 8.0, with 5 mmol/L calcium acetate and stored at 4°C. Then added 2 μL stock solution to tail tip samples that had been previously placed into 200 μL GNT-K buffer (10 mmol/L Tris-HCl, pH 8.5, 50 mmol/L KCl, 1.5 mmol/L MgCl_2_, 0.01% gelatin, 0.45% Nonidet P-40, and 0.45% Tween20). The samples were digested for no more than 2 h at 56°C on a dry heat block. The samples were then heated at 95°C for 15 min to inactivate proteinase K and were briefly centrifuged at 13,000 g for 1 min.

Immediately after centrifugation, 2 μL DNA containing supernatant was transferred to 20-μL reactions. 2× Taq Master Mix (Vayzme, P112-01) was used for the PCR reactions. The amplified products were then used for DNA electrophoresis. Primer pairs for genotyping are shown in Table S2.

### Isolation of mitochondria from hearts

Mitochondria were isolated from mouse heart tissues as previously described (Wieckowski, Giorgi, Lebiedzinska, Duszynski, & Pinton, 2009). In brief, mice were euthanized and heart tissue was immediately washed in ice-cold PBS (three times with PBS to remove the blood), then tissue was cut into small pieces using scissors. The used PBS solution was discarded and the tissue was washed once again with 10 mL of fresh, ice-cold IB-1. The heart tissue was transferred to an ice-cold glass/Teflon Potter Elvehjem homogenizer. IB-1 was added in the ratio of 4 mL of buffer per gram of heart tissue. Homogenization, as well as the following steps, were carried out at 4°C to minimize the activation of proteases and phospholipases. After homogenization was completed, tissue was transferred to a centrifuge and spun at 700 g for 10 min at 4°C (repeated 3 times). The supernatant was collected and centrifuged at 9,000 g for 10 min at 4°C. The supernatant (representing the cytosolic fraction containing lysosomes and microsomes) was then discarded, and the pellet (containing the mitochondria) was gently resuspended in 20 mL of ice-cold IB-2. The mitochondrial suspension was then centrifuged at 10,000 g for 10 min at 4°C (repeated 2 times). The pellet representing the crude mitochondrial fraction was used for further experimentation.

### Adult mouse cardiomyocyte isolation

Adult mouse cardiomyocytes were isolated according to a previously published method (Li et al., 2021). Briefly, after sacrifice, the mouse heart was immediately harvested and cannulated. The heart was then perfused with the modified Krebs-Ringer buffer (HEPES 20 mM, NaCl 137 mmol/L, KCl 5.4 mmol/L, MgCl_2_·6H_2_O 1.2 mmol/L, Na_2_HPO_4_ 1.2 mmol/L, glucose 10 mmol/L, taurin 10 mmol/L, pH 7.4) with collagenase II (0.8 mg/mL, Worthington, LS004177) and protease (0.04 mg/mL; Sigma-Aldrich, P5147). After digestion for approximately 25 min, atria were removed and ventricles were dissociated by pipetting. After filtration by cell strainer, adult cardiomyocytes were separated and enriched by sedimentation twice, and stored for further use.

### Seahorse metabolic assays

The OCR was measured using a Seahorse XFe24 analyzer (Agilent, USA). NRCMs were seeded at 50,000 cells/well on to XFe24 microplates (Agilent, 100777) in high glucose DMEM medium (Hyclone) containing 10% FBS, penicillin and streptomycin at 37[ and 5% CO_2_. After 48 h, cells were treated with lentivirus or transfected siRNA, and after 72 h, seahorse assay medium was prepared by adding pyruvate, glutamine and glucose to XF Base Medium. Cells were then cultured in the assay medium for 1 h in a 37[ and non-CO_2_ incubator prior to the assay. 50 μg of mitochondrial protein from the hearts was added to XFe24 microplates. A mitochondrial stress test kit (Agilent, 103015-100) was used to monitor OCR where baseline measurements were made followed by sequential injection of ADP (4 mmol/L), oligomycin (4 μmol/L), FCCP (2 μmol/L), and Rotenone (1 μmol/L). Data were analyzed in seahorse wave software. The ECAR was measured using a Seahorse XFe24 analyzer (Seahorse Bioscience) following the manufacturer’s protocol. NRCMs were cultured and treated before assay following a similar protocol as in mitochondrial stress test assay. A glycolysis stress test kit (Agilent, 103020-100) was used to monitor ECAR where baseline measurements were made followed by sequential injection of glucose (10 mmol/L), oligomycin (1 μmol/L), 2-DG (50 mmol/L). Data were analyzed in Seahorse wave software.

### Hypoxia and reoxygenation assay

NRCMs were cultured for 48h under normoxia before being transferred to a hypoxia incubator with a gas mixture containing 37[, 5% CO_2_ and 0.1% O_2_ balanced with nitrogen. After 12 h of hypoxic culture, the cells were reoxygenated in the normal incubator containing 37[, 5% CO_2_ for 1 h. Cells were immediately harvested at the indicated time points.

### Western blot analysis

Cells and left ventricular tissue were lysed with RIPA lysis buffer (Beyotime, P0013B, Shanghai, China) and protein concentrations were measured using the BCA assay kit (Beyotime, P0010). The total proteins were separated by sodium dodecyl sulfate-polyacrylamide gel electrophoresis and transferred to nitrocellulose membranes. After blocking, the membranes were incubated with anti-AARS2 (1:1000, Abcam, ab197367, Cambridge, UK), anti-B-cell lymphoma-2 (Bcl-2, 1:1000, CST, 3498S), anti-Bcl-2 associated X (BAX, 1:1000, CST, 14796S), anti-PKM2 (1:1000, CST, 4053S), anti-PKM1 (1:1000, Proteintech, 15821-1-AP), anti-PDK4 (1:1000, Bioss, bs-0682R) and anti-LDHA (1:1000, Bioss, bs-34202R). After washing with TBS with Tween-20, Goat Anti-Rabbit IgG (H&L)-HRP Conjugated (1:10000, EASYBIO, BE0101-100) was added, HRP-conjugated GAPDH Monoclonal antibody (1:10000, Proteintech, HRP-60004), anti-β-actin (1:10000, Proteintech, HRP-60008) and anti-α-Tubulin (1:10000, Proteintech, HRP-66031) were used as a control for normalization. Protein signals were detected with Super ECL Detection Reagent (Yeasen, 36208ES60) in a ChemiDoc™ MP Imaging System (Bio-Rad). Native proteins were extracted using Column Tissue & Cell Protein Extraction Kit (Epizyme, PC201 plus) according to the manufacturer’s instructions.

### Co-Immunoprecipitation

For immunoprecipitation, cells with an 85% confluent at 15-cm plate were used and equilibrated to serum free DMEM for 2 h prior to immunoprecipitation. The total protein from cell lysate was immunoprecipitated. Firstly, the extract was incubated with anti-PKM2 for 24 h at 4°C. The protein A/G Agarose (Beyotime, P2012) was added and further incubated for 3 h at 4°C followed by centrifuged at 12,000 g for 5 min. Then recovered the precipitate for washing, resuspended it in 40 μL SDS lysis buffer and boiled for 5 min, and finally analyzed the precipitate by immunoblotting with the indicated antibody.

### Immunofluorescence cytochemistry

Heart tissues were embedded in OCT compound and sectioned at 6 µm on a cryostat. After fixation in PFA, sections were permeabilized with 1% Triton X-100 for 10 min and blocked with 1% BSA for 60 min. NRCMs on cell culture plates were fixed in 4% PFA for 30 min and permeabilized with 1% Triton X-100 for 10 min and blocked with 1% BSA for 1 h. Then, slices and NRCMs were incubated with rabbit anti-AARS2 (1:300; Abcam, ab197367), rabbit anti-Ki67 (1:300; CST, 12075S), rabbit anti-pH3 (1:300; CST, 53348S), rabbit anti-CD31 (1:300; Abcam, ab182981), mouse Anti-Cardiac Troponin T (cTnT, 1:300; Abcam, ab8295) and mouse anti-α-SMA (1:300; CST, 48938S) at 4°C overnight. After washing with 1% PBS-Tween, Alexa Fluor 488-conjugated anti-rabbit and Alexa Fluor 555-conjugated anti-mouse secondary antibodies (1:500; Abcam, ab150077, ab150118) were added.

Terminal deoxynucleotidyl transferase dUTP nick end labeling (TUNEL) assay was performed using the TUNEL Apoptosis Detection Kit (Alexa Fluor 488; Beyotime, C1088) according to the manufacturer’s instructions. MitoSOX (Thermo Fisher Scientific, M36008) staining was applied directly on the NRCMs for 10 min following the manufacturer’s instructions. All the sections were counterstained with 4’, 6-diamidino-2-phenylindole (DAPI, Abcam, ab104139). Fluorescent cells and tissues were visualized and digital images were captured using Axio Scan Z1 (Carl Zeiss, Germany).

### WGA staining

To determine the cross-section areas of CMs, heart sections were dewaxed in xylene, rehydrated in an ethanol series, washed in PBS, and then incubated for 1 h with Wheat Germ Agglutinin, Alexa Fluor 488 Conjugate (5 μg/mL, Invitrogen, W11261). Added mouse Anti-cTnT (1:300; Abcam, ab8295) at 4°C overnight. Then added Alexa Fluor 555-conjugated anti-mouse secondary antibodies (1:500; Abcam, ab150118) for 1 h. The slides were then rinsed in PBS and co-stained with DAPI. To quantify the size of cells, images at 20 × magnification were analyzed using ImageJ.

### LDH release assay

The mice serum and cell lysate were collected and analyzed for LDH release using an LDH-release kit (Beyotime, C0016) following the manufacturer’s instructions.

### Quantitative Real-Time PCR

mRNA was isolated from cells or tissues with a Total RNA kit (Tiangen, DP424, Shanghai, China), and reverse-transcribed with an RT Master Mix kit (Yeasen, 111141ES60, Shanghai, China). Quantitative Real-Time PCR (qRT-PCR) was performed with the SYBR Premix Ex Taq kit (Yeasen, 11184ES08) on an AB 7500 Fast Real-Time PCR System (Applied Biosystems). β-actin and GAPDH were used as an internal control. The sequences of primers are summarized in Table S3. The fold changes in mRNA expression levels were normalized to β[actin and GAPDH using the ^ΔΔ^Ct method.

### Metabolite Profiling

Frozen mouse heart tissues were homogenized with TissueLyser II by adding the −20°C-cold high-performance liquid chromatography-grade methanol to the final concentration of 100 mg/mL. The denatured protein was pelleted by centrifuging the tube at 14,000 rpm for 20 min at 4[ and the supernatant of individual sample was transferred into the new tube for metabolite profiling analysis. Before the LC-MS based analysis, 10 μL of supernatant was diluted with 90 μL of methanol for positive mode analysis, and 30 μL of supernatant was diluted with 70 μL of acetonitrile/methanol (3:1) for negative mode analysis, respectively.

10 μL of reconstituted sample was loaded onto either a 150 × 2.1 mm Atlantis HILIC column (Waters, Milford, MA) for positive mode analysis or 5 μL was loaded on a 100 × 2.1 mm 3.5 μm XBridge amide column (Waters) for negative mode analysis using an HTS PAL Autosampler (Leap Technologies, Carrboro, NC) or Agilent 1290 Infinity Autosampler. The metabolites were separated using an Agilent 1200 Series high-performance liquid chromatography system (Agilent Technologies, Santa Clara, CA) coupled to a 4000-QTRAP mass spectrometer (AB SCIEX, Foster City, CA) in positive mode analysis; an Agilent 1290 infinity high-performance liquid chromatography binary pump system (Agilent Technologies) coupled to a 6490-QQQ mass spectrometer (Agilent Technologies) in negative mode analysis MultiQuant software v2.1 (AB SCIEX) and MassHunter quantitative software were used for automated peak integration, respectively, and metabolite peaks were also assessed manually for quality of peak integration.

### Ribosome Profiling Sequencing

Ribo-Seq of NRCMs was performed by Novogene. Briefly, use RNase I to digest the unprotected RNA, leaving only the ribosome-protected mRNA fragments, and SE50. Differential gene expression of control and AARS2^OE^ groups was analyzed using the DESeq2 R package. DESeq2 provides statistical routines for determining digital differential gene expression data using a model based on the negative binomial distribution. We used the cluster Profiler R package to test the statistical enrichment of differentially expressed genes.

### Statistical analysis

Values are reported as mean ± s.e.m. unless indicated otherwise. The 2-tailed non-parametric Student’s T-test was used for 2-group comparisons by Prism 9.0 (GraphPad Software, San Diego, CA, USA) statistical software. In addition, one-way analysis of variance analysis (ANOVA) was used to evaluate the statistical significance for multiple-group comparisons. Values of P<0.05 were considered statistically significant.

## Acknowledgements

The authors thank National Center for Protein Sciences at Peking University in Beijing; and the members of Dr. Jing-Wei Xiong’s laboratory for helpful discussions and technical assistance.

## Data availability

RNA-Seq data have been deposited at ArrayExpress (E-MTAB-13767).

## Author contribution

ZW Zhang designed and performed experiments, analyzed data, and wrote the manuscript; LX Zheng, Y Chen, YY Chen and JJ Hou performed the experiments and analyzed the data. CL Xiao and XJ Zhu analyzed the data and managed the project. SM Zhao provided AARS2^fl/fl^ and AARS2^Tg^ mouse lines, designed the experiments and analyzed the data. JW Xiong conceived and designed this work, analyzed data, and wrote the manuscript. All the authors approved the final draft for submission.

---

**Figure 1–figure supplement 1.**
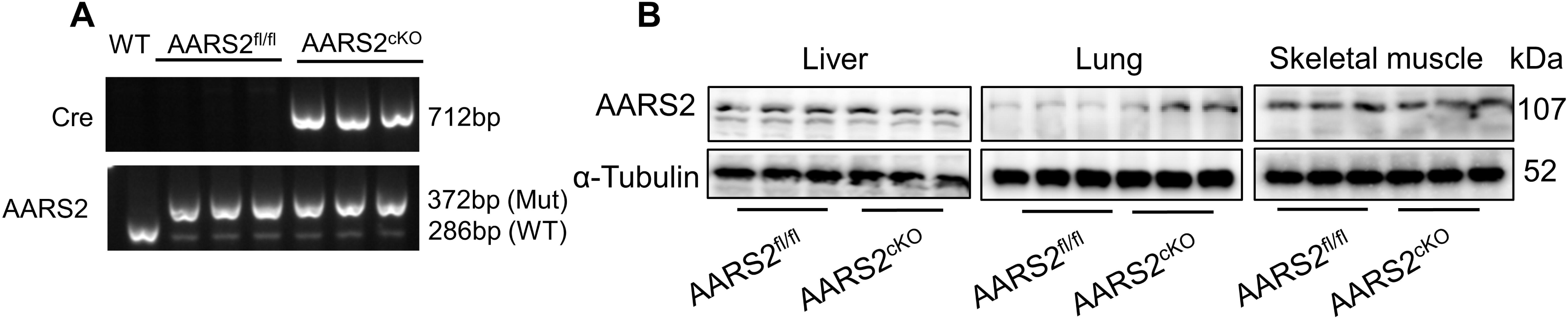
Genotyping of cardiomyocyte-specific AARS2 knockout mice. (A) Genotyping of AARS2^fl/fl^ and AARS2^cKO^ mice (n = 3; Mut, mutant). (B) Western Blot showing normal expression of AARS2 proteins in the liver, lung, and skeletal muscle of AARS2^fl/fl^ and AARS2^cKO^ mice (n = 3).

**Figure 2–figure supplement 1.**
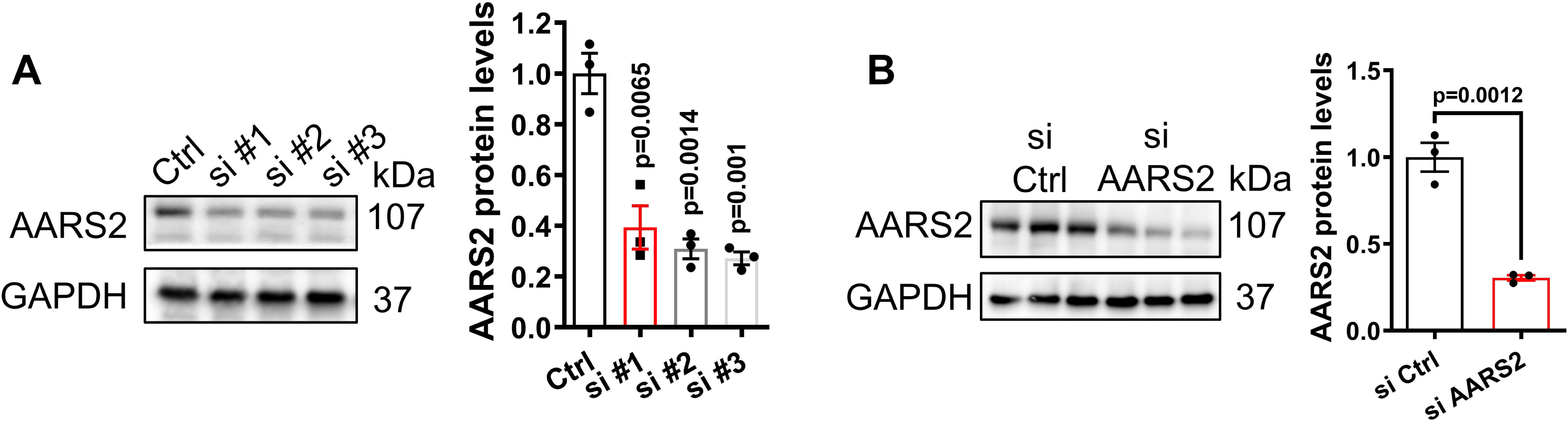
Evaluating AARS2 siRNA for knockout efficiency of AARS2 proteins in NRCMs. (A) Western blots showing knockdown efficiency of 3 different AARS2 siRNAs in NRCMs, respectively (n = 3). (B) Western blots confirming knockdown efficiency of AARS2 siRNA#3 in NRCMs (n = 3). Ctrl, control; si, small interfering; mean ± s.e.m.

**Figure 3–figure supplement 1.**
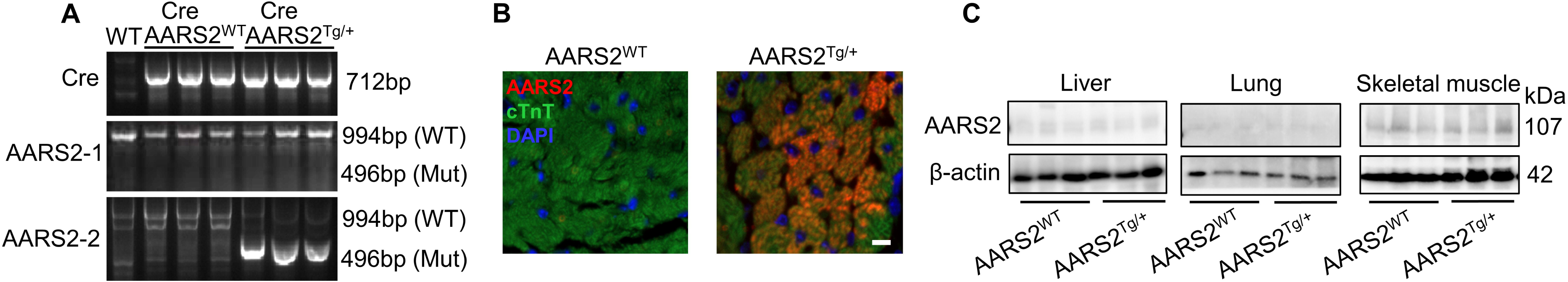
Cardiomyocyte-specific overexpression of AARS2 in the heart but not in the liver, lung, and skeletal muscle. (A) Genotyping of wild type (WT), α-MHC-MerCreMer; AARS2^WT^ (AARS2^WT^) control, and α-MHC-MerCreMer; AARS2^Tg/+^ (AARS2^Tg/+^) mice (n = 3). (B) Immunofluorescence staining showing ectopic AARS2 proteins in cardiomyocytes of AARS2^Tg/+^ transgenic mice compared with AARS2^WT^ control mice (scale bar, 50 μm). (C) Western blots showing comparable expression of AARS2 proteins in the liver, lung, and skeletal muscle of AARS2^WT^ control and AARS2^Tg/+^ transgenic mice (n = 3). Mean ± s.e.m.

**Figure 3–figure supplement 2.**
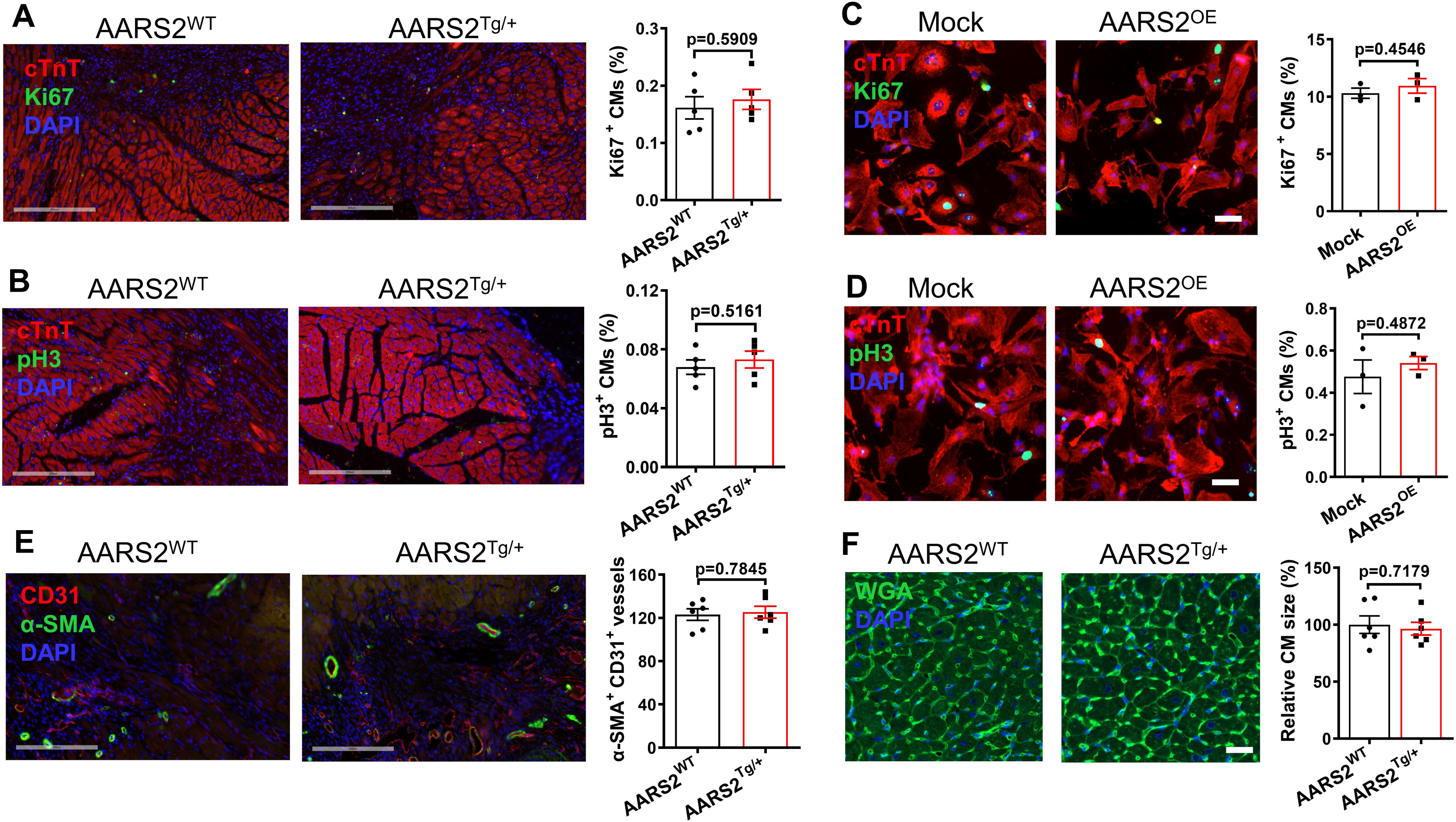
Overexpression of AARS2 in cardiomyocytes has no apparent effect on cardiomyocyte proliferation, hypertrophy, and angiogenesis after MI. (A–B) At 7 days after MI, immunofluorescence staining showing comparable cTnT^+^/Ki67^+^ (A) and cTnT^+^/pH3^+^ (B) cardiomyocytes in the heart sections of AARS2^WT^ and AARS2^Tg/+^ groups (scale, 200 μm; n = 5). (C–D) Immunofluorescence staining showing comparable numbers of cTnT^+^/Ki67^+^ (C) and cTnT^+^/pH3^+^ (D) cardiomyocytes at 48 h after infection of NRCMs with Mock lentivirus or AARS2 lentivirus (scale, 50 μm; n = 3). (E) At 7 days after MI, immunofluorescence staining showing comparable CD31^+^/α-SMA^+^ coronary vessels in the heart sections of AARS2^WT^ and AARS2^Tg/+^ groups (scale, 200 μm; n = 5). (F) WGA staining showing no evident cardiomyocyte hypertrophy in heart slices of AARS2^WT^ and AARS2^Tg/+^ groups at 28 days after MI (scale, 100 μm; n = 6). Mean ± s.e.m.

**Figure 6–figure supplement 1.**
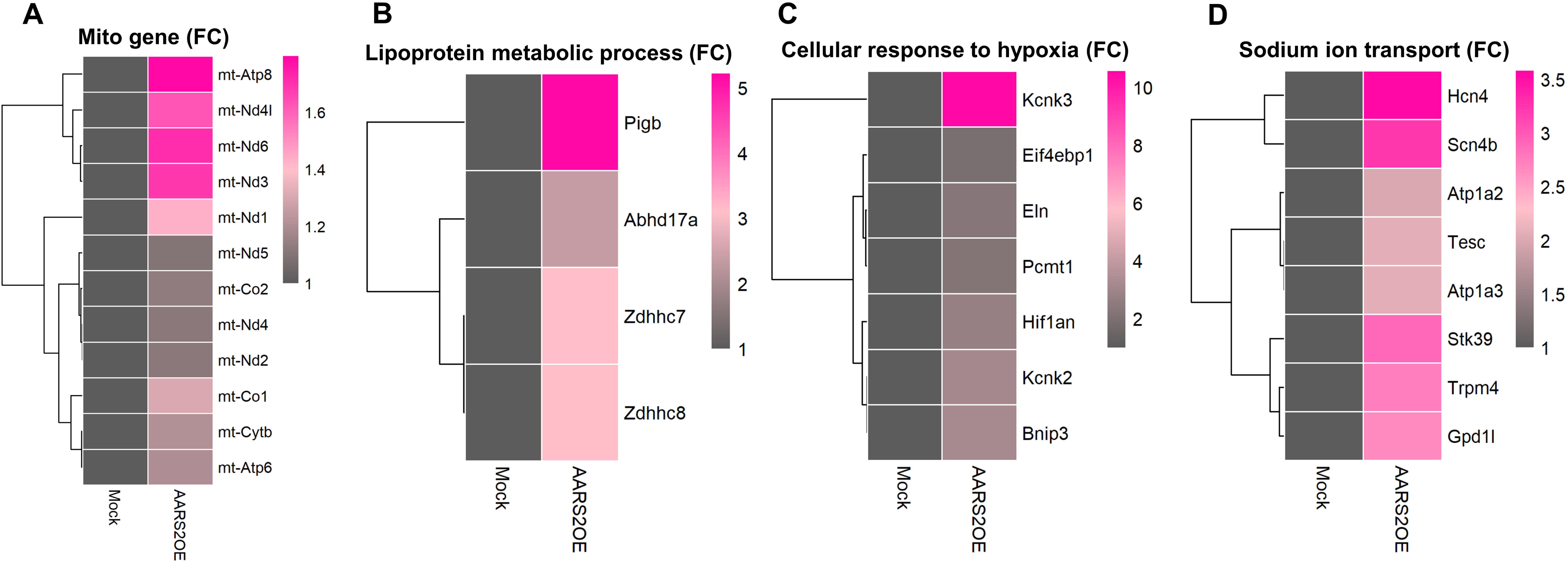
Overexpression of AARS2 increases the translation level of some cellular proteins. (A–D) Ribosome RNA-Seq showing elevated translation of signaling pathways of genes encoded by mitochondria(A), lipoprotein metabolic process (B), cellular response to hypoxia (C), and sodium ion transport (D) in the AARS2^OE^ NRCMs compared to the Mock NRCMs. FC, fold changes.

## References

1. Chen, L., Shi, Y., Liu, S., Cao, Y., Wang, X., & Tao, Y. (2014). PKM2: the thread linking energy metabolism reprogramming with epigenetics in cancer. Int J Mol Sci, 15(7), 11435–11445. doi:10.3390/ijms150711435

2. Chuang, H. M., Su, H. L., Li, C., Lin, S. Z., Yen, S. Y., Huang, M. H., . . . Harn, H. J. (2016). The Role of Butylidenephthalide in Targeting the Microenvironment Which Contributes to Liver Fibrosis Amelioration. Front Pharmacol, 7, 112. doi:10.3389/fphar.2016.00112

3. Dallabona, C., Diodato, D., Kevelam, S. H., Haack, T. B., Wong, L. J., Salomons, G. S., . . . van der Knaap, M. S. (2014). Novel (ovario) leukodystrophy related to AARS2 mutations. Neurology, 82(23), 2063–2071. doi:10.1212/wnl.0000000000000497

4. Dayton, T. L., Gocheva, V., Miller, K. M., Israelsen, W. J., Bhutkar, A., Clish, C. B., . . . Vander Heiden, M. G. (2016). Germline loss of PKM2 promotes metabolic distress and hepatocellular carcinoma. Genes Dev, 30(9), 1020–1033. doi:10.1101/gad.278549.116

5. Dayton, T. L., Jacks, T., & Vander Heiden, M. G. (2016). PKM2, cancer metabolism, and the road ahead. EMBO Rep, 17(12), 1721–1730. doi:10.15252/embr.201643300

6. Du, J., Zheng, L., Gao, P., Yang, H., Yang, W. J., Guo, F., . . . Xiong, J. W. (2022). A small-molecule cocktail promotes mammalian cardiomyocyte proliferation and heart regeneration. Cell Stem Cell, 29(4), 545–558.e513. doi:10.1016/j.stem.2022.03.009

7. Euro, L., Konovalova, S., Asin-Cayuela, J., Tulinius, M., Griffin, H., Horvath, R., Tyynismaa, H. (2015). Structural modeling of tissue-specific mitochondrial alanyl-tRNA synthetase (AARS2) defects predicts differential effects on aminoacylation. Front Genet, 6, 21. doi:10.3389/fgene.2015.00021

8. Fine, A. S., Nemeth, C. L., Kaufman, M. L., & Fatemi, A. (2019). Mitochondrial aminoacyl-tRNA synthetase disorders: an emerging group of developmental disorders of myelination. J Neurodev Disord, 11(1), 29. doi:10.1186/s11689-019-9292-y

9. Frangogiannis, N. G. (2015). Pathophysiology of Myocardial Infarction. Compr Physiol, 5(4), 1841–1875. doi:10.1002/cphy.c150006

10. Gorman, G. S., Chinnery, P. F., DiMauro, S., Hirano, M., Koga, Y., McFarland, R., . . . Turnbull, D. M. (2016). Mitochondrial diseases. Nat Rev Dis Primers, 2, 16080. doi:10.1038/nrdp.2016.80

11. Ikeda, Y., Tanaka, T., & Noguchi, T. (1997). Conversion of non-allosteric pyruvate kinase isozyme into an allosteric enzyme by a single amino acid substitution. J Biol Chem, 272(33), 20495–20501. doi:10.1074/jbc.272.33.20495

12. Israelsen, W. J., & Vander Heiden, M. G. (2015). Pyruvate kinase: Function, regulation and role in cancer. Semin Cell Dev Biol, 43, 43–51. doi:10.1016/j.semcdb.2015.08.004

13. Ju, J., Zhang, H., Lin, M., Yan, Z., An, L., Cao, Z., . . . Zhou, Z. (2024). The alanyl-tRNA synthetase AARS1 moonlights as a lactyl-transferase to promote YAP signaling in gastric cancer. J Clin Invest. doi:10.1172/jci174587

14. Kalogeris, T., Baines, C. P., Krenz, M., & Korthuis, R. J. (2012). Cell biology of ischemia/reperfusion injury. Int Rev Cell Mol Biol, 298, 229–317. doi:10.1016/b978-0-12-394309-5.00006-7

15. Li, Q., Li, C., Elnwasany, A., Sharma, G., An, Y. A., Zhang, G., . . . Wang, Z. V. (2021). PKM1 Exerts Critical Roles in Cardiac Remodeling Under Pressure Overload in the Heart. Circulation, 144(9), 712–727. doi:10.1161/circulationaha.121.054885

16. Magadum, A., Singh, N., Kurian, A. A., Munir, I., Mehmood, T., Brown, K., . . . Zangi, L. (2020). Pkm2 Regulates Cardiomyocyte Cell Cycle and Promotes Cardiac Regeneration. Circulation, 141(15), 1249–1265. doi:10.1161/circulationaha.119.043067

17. Mao, Y., Zhang, J., Zhou, Q., He, X., Zheng, Z., Wei, Y., . . . Zhao, S. (2024). Hypoxia induces mitochondrial protein lactylation to limit oxidative phosphorylation. Cell Res, 34(1), 13–30. doi:10.1038/s41422-023-00864-6

18. Méndez-Lucas, A., Li, X., Hu, J., Che, L., Song, X., Jia, J., . . . Chen, X. (2017). Glucose Catabolism in Liver Tumors Induced by c-MYC Can Be Sustained by Various PKM1/PKM2 Ratios and Pyruvate Kinase Activities. Cancer Res, 77(16), 4355–4364. doi:10.1158/0008-5472.Can-17-0498

19. Mendis, S., Davis, S., & Norrving, B. (2015). Organizational update: the world health organization global status report on noncommunicable diseases 2014; one more landmark step in the combat against stroke and vascular disease. Stroke, 46(5), e121–122. doi:10.1161/strokeaha.115.008097

20. Meyer-Schuman, R., & Antonellis, A. (2017). Emerging mechanisms of aminoacyl-tRNA synthetase mutations in recessive and dominant human disease. Hum Mol Genet, 26(R2), R114–r127. doi:10.1093/hmg/ddx231

21. Ni, L., Lin, B., Hu, L., Zhang, R., Fu, F., Shen, M., . . . Shi, D. (2022). Pyruvate Kinase M2 Protects Heart from Pressure Overload-Induced Heart Failure by Phosphorylating RAC1. J Am Heart Assoc, 11(11), e024854. doi:10.1161/jaha.121.024854

22. Nielsen, S. K., Hansen, F., Schrøder, H. D., Wibrand, F., Gustafsson, F., & Mogensen, J. (2020). Recessive Inheritance of a Rare Variant in the Nuclear Mitochondrial Gene for AARS2 in Late-Onset Dilated Cardiomyopathy. Circ Genom Precis Med, 13(5), 560–562. doi:10.1161/circgen.120.003086

23. Ognjenović, J., & Simonović, M. (2018). Human aminoacyl-tRNA synthetases in diseases of the nervous system. RNA Biol, 15(4-5), 623–634. doi:10.1080/15476286.2017.1330245

24. Palsson-McDermott, E. M., Curtis, A. M., Goel, G., Lauterbach, M. A. R., Sheedy, F. J., Gleeson, L. E., . . . O’Neill, L. A. J. (2015). Pyruvate Kinase M2 Regulates Hif-1α Activity and IL-1β Induction and Is a Critical Determinant of the Warburg Effect in LPS-Activated Macrophages. Cell Metab, 21(2), 347. doi:10.1016/j.cmet.2015.01.017

25. Pearce, S., Nezich, C. L., & Spinazzola, A. (2013). Mitochondrial diseases: translation matters. Mol Cell Neurosci, 55, 1–12. doi:10.1016/j.mcn.2012.08.013

26. Penna, C., Mancardi, D., Rastaldo, R., & Pagliaro, P. (2009). Cardioprotection: a radical view Free radicals in pre and postconditioning. Biochim Biophys Acta, 1787(7), 781–793. doi:10.1016/j.bbabio.2009.02.008

27. Prakasam, G., Iqbal, M. A., Bamezai, R. N. K., & Mazurek, S. (2018). Posttranslational Modifications of Pyruvate Kinase M2: Tweaks that Benefit Cancer. Front Oncol, 8, 22. doi:10.3389/fonc.2018.00022

28. Saraste, M. (1999). Oxidative phosphorylation at the fin de siècle. Science, 283(5407), 1488–1493. doi:10.1126/science.283.5407.1488

29. Schieber, M., & Chandel, N. S. (2014). ROS function in redox signaling and oxidative stress. Curr Biol, 24(10), R453–462. doi:10.1016/j.cub.2014.03.034

30. Sissler, M., González-Serrano, L. E., & Westhof, E. (2017). Recent Advances in Mitochondrial Aminoacyl-tRNA Synthetases and Disease. Trends Mol Med, 23(8), 693–708. doi:10.1016/j.molmed.2017.06.002

31. Sohal, D. S., Nghiem, M., Crackower, M. A., Witt, S. A., Kimball, T. R., Tymitz, K. M., . . . Molkentin, J. D. (2001). Temporally regulated and tissue-specific gene manipulations in the adult and embryonic heart using a tamoxifen-inducible Cre protein. Circ Res, 89(1), 20–25. doi:10.1161/hh1301.092687

32. Strait, J. B., & Lakatta, E. G. (2012). Aging-associated cardiovascular changes and their relationship to heart failure. Heart Fail Clin, 8(1), 143–164. doi:10.1016/j.hfc.2011.08.011

33. Tang, Y., Feng, M., Su, Y., Ma, T., Zhang, H., Wu, H., . . . Xu, D. (2023). Jmjd4 Facilitates Pkm2 Degradation in Cardiomyocytes and Is Protective Against Dilated Cardiomyopathy. Circulation, 147(22), 1684–1704. doi:10.1161/circulationaha.123.064121

34. Thygesen, K., Alpert, J. S., Jaffe, A. S., Chaitman, B. R., Bax, J. J., Morrow, D. A., & White, H. D. (2018). Fourth Universal Definition of Myocardial Infarction (2018). J Am Coll Cardiol, 72(18), 2231–2264. doi:10.1016/j.jacc.2018.08.1038

35. Vander Heiden, M. G., Cantley, L. C., & Thompson, C. B. (2009). Understanding the Warburg effect: the metabolic requirements of cell proliferation. Science, 324(5930), 1029–1033. doi:10.1126/science.1160809

36. Vasilescu, C., Ojala, T. H., Brilhante, V., Ojanen, S., Hinterding, H. M., Palin, E., . . . Suomalainen, A. (2018). Genetic Basis of Severe Childhood-Onset Cardiomyopathies. J Am Coll Cardiol, 72(19), 2324–2338. doi:10.1016/j.jacc.2018.08.2171

37. Wei, Y., Miao, Q., Zhang, Q., Mao, S., Li, M., Xu, X., . . . Hu, G. (2023). Aerobic glycolysis is the predominant means of glucose metabolism in neuronal somata, which protects against oxidative damage. Nat Neurosci, 26(12), 2081–2089. doi:10.1038/s41593-023-01476-4

38. Whelan, R. S., Kaplinskiy, V., & Kitsis, R. N. (2010). Cell death in the pathogenesis of heart disease: mechanisms and significance. Annu Rev Physiol, 72, 19–44. doi:10.1146/annurev.physiol.010908.163111

39. Wieckowski, M. R., Giorgi, C., Lebiedzinska, M., Duszynski, J., & Pinton, P. (2009). Isolation of mitochondria-associated membranes and mitochondria from animal tissues and cells. Nat Protoc, 4(11), 1582–1590. doi:10.1038/nprot.2009.151

40. Wong, N., Ojo, D., Yan, J., & Tang, D. (2015). PKM2 contributes to cancer metabolism. Cancer Lett, 356(2 Pt A), 184-191. doi:10.1016/j.canlet.2014.01.031

41. Wu, X., Liu, L., Zheng, Q., Hao, H., Ye, H., Li, P., & Yang, H. (2021). Protocatechuic aldehyde protects cardiomycoytes against ischemic injury via regulation of nuclear pyruvate kinase M2. Acta Pharm Sin B, 11(11), 3553–3566. doi:10.1016/j.apsb.2021.03.021

42. Zhang, Q., Wang, L., Wang, S., Cheng, H., Xu, L., Pei, G., Wei, Q. (2022). Signaling pathways and targeted therapy for myocardial infarction. Signal Transduct Target Ther, 7(1), 78. doi:10.1038/s41392-022-00925-z

43. Zhang, T., Zhang, Y., Cui, M., Jin, L., Wang, Y., Lv, F., Xiao, R. P. (2016). CaMKII is a RIP3 substrate mediating ischemia- and oxidative stress-induced myocardial necroptosis. Nat Med, 22(2), 175–182. doi:10.1038/nm.4017

44. Zhang, X., Li, J., Zhang, Y., Gao, M., Peng, T., & Tian, T. (2022). AARS2-Related Leukodystrophy: a Case Report and Literature Review. Cerebellum. doi:10.1007/s12311-022-01369-5

45. Zhang, Z., Chen, Y., Zheng, L., Du, J., Wei, S., Zhu, X., & Xiong, J. W. (2023). A DUSP6 inhibitor suppresses inflammatory cardiac remodeling and improves heart function after myocardial infarction. Dis Model Mech, 16(5). doi:10.1242/dmm.049662

46. Zhao, X., Han, J., Zhu, L., Xiao, Y., Wang, C., Hong, F., . . . Guan, M. X. (2018). Overexpression of human mitochondrial alanyl-tRNA synthetase suppresses biochemical defects of the mt-tRNA(Ala) mutation in cybrids. Int J Biol Sci, 14(11), 1437–1444. doi:10.7150/ijbs.27043

47. Zhou, Y., Chen, B., Li, L., Pan, H., Liu, B., Li, T., . . . Cao, Y. (2019). Novel alanyl-tRNA synthetase 2 (AARS2) homozygous mutation in a consanguineous Chinese family with premature ovarian insufficiency. Fertil Steril, 112(3), 569–576.e562. doi:10.1016/j.fertnstert.2019.05.005

48. Zong, Z., Xie, F., Wang, S., Wu, X., Zhang, Z., Yang, B., & Zhou, F. (2024). Alanyl-tRNA synthetase, AARS1, is a lactate sensor and lactyltransferase that lactylates p53 and contributes to tumorigenesis. Cell. doi:10.1016/j.cell.2024.04.002

