## Supplementary table S1-S3 for "AARS2 ameliorates myocardial ischemia *via* fine-tuning PKM2-mediated metabolism"

**Supplementary Data**

**Table S1. AARS2 siRNA** **sequences**

| Genes |  | Primers | Sequence (5' to 3') |
| --- | --- | --- | --- |
| AARS2-si-#1 |  | Forward | GCUUCCGACGAGUAGCUAACATT |
|  |  | Reverse | UGUUAGCUACUCGUCGGAAGCTT |
| AARS2-si-#2 |  | Forward | CGUUCGGACUGCAAGAGAACUTT |
|  |  | Reverse | AGUUCUCUUGCAGUCCGAACGTT |
| AARS2-si-#3 |  | Forward | GGAGGAGCUGCACCGUCAAGGTT |
|  |  | Reverse | CCUUGACGGUGCAGCUCCUCCTT |

**Table S2. Table S2. Genotyping primer sequences**

AARS2 transgenic mouse primers

| Primer | Sequences (5'-3') |
| --- | --- |
| P1 Forward | TCAGATTCTTTTATAGGGGACACA |
| P1 Reverse  P2 Forward | TAAAGGCCACTCAATGCTCACTAA  TGCCCTCAGTATAGCCCAAACC |
| P2 Reverse | GCAGCCAAGGAAAGGACGATGATT |

The molecular weight of the two pairs of primers was the same, generating a WT band: 994bp; and Mut (mutant) band: 496bp.

AARS2 knockout mouse primers

| Primers | Sequences (5' to 3') |
| --- | --- |
| P1 Forward | AAGCAACAGGAGAAGAGGTGTTGG |
| P1 Reverse | TAACCATCTCAGCAGCCCAGCAT |

The pair of primers generating a WT band: 268bp, and MUT band: 372bp.

Cre primers

| Primers | Sequences (5' to 3') |
| --- | --- |
| P1 Forward | AATGCTTCTGTCCGTTTGC |
| P1 Reverse | ACCAGAGTCATCCTTAGCG |

Cre primer pair generating a 712-bp band.

**Table S3. Primers used for real-time PCR analysis**

| Genes | Primers | Sequences (5' to 3') |
| --- | --- | --- |
| rat PKM2 | Forward | GATCTGAAGTACGCCCGAGG |
| rat PKM2 | Reverse | GAATGAAGGCAGTCCCTGCT |
| rat β-actin | Forward | AACCTTCTTGCAGCTCCTCC |
| rat β-actin | Reverse | TACCCACCATCACACCCTGG |
| mouse β-actin | Forward | GGCTGTATTCCCCTCCATCG |
| mouse β-actin | Reverse | CCAGTTGGTAACAATGCCATGT |
| mouse AARS2 | Forward | AAGTTACGGTATGCTGAGCCG |
| mouse AARS2 | Reverse | AACTACACGTCGGAAGCCTG |
| mouse PKM2 | Forward | TGTCTGGAGAAACAGCCAAG |
| mouse PKM2 | Reverse | TCCTCGAATAGCTGCAAGTG |
